# Perception of naturally dead conspecifics impairs health and longevity through serotonin signaling in *Drosophila*

**DOI:** 10.1101/515312

**Authors:** Tuhin S. Chakraborty, Christi M. Gendron, Yang Lyu, Allyson S. Munneke, Madeline N. DeMarco, Zachary W. Hoisington, Scott D. Pletcher

## Abstract

Sensory perception modulates health and aging across taxa. Understanding the nature of relevant cues and the mechanisms underlying their action may lead to novel interventions that improve the length and quality of life. In humans, psychological trauma is often associated with the recognition of dead individuals, with chronic exposure leading to persistent mental health issues including depression and post-traumatic stress disorder. The mechanisms that link mental and physical health, and the degree to which these are shared across species, remain largely unknown. Here we show that the vinegar fly, *Drosophila melanogaster*, has the capability to perceive dead conspecifics in its environment and that this perceptive experience induces both short- and long-term effects on health and longevity. Death perception is mediated by visual and olfactory cues, and remarkably, its effects on aging are eliminated by targeted attenuation of serotonin signaling. Our results suggest a complex perceptive ability in *Drosophila* that reveals deeply conserved mechanistic links between psychological state and aging, the roots of which might be unearthed using invertebrate model systems.

---

In humans, psychological stress is often induced by our perception of alarming or distasteful stimuli, and the recognition of dead individuals is one of the most potent. This perceptive ability is not uncommon in the animal kingdom, as individuals from a wide range of species respond to their dead. Social insects, including ants and honey bees, exhibit necrophoresis in which dead colony members are systematically removed from the nest to promote hygienic conditions ^1^. Dead zebrafish scents provoke defensive behavior in live individuals ^2^, and the sight of a dead conspecific induces alarm calling in scrub-jays ^3^, suggesting that dead individuals may indicate danger. Elephants and nonhuman primates exhibit stereotypical behaviors toward dead individuals associated with permanent loss of a group member ^4,5^. Such natural behaviors resemble human emotions that are associated with our own experiences of death, which include anger, fear, anxiety, and depression. Short-term effects of these psychologically stressful conditions include emotional dysregulation, hyperactivity, eating disorders, and alcohol abuse ^6^. Long-term consequences of exposure to death are evident in first responders as well as active military troops; they include both psychological effects, such as post-traumatic stress disorder and depression, and physical effects, such as headaches, fatigue, and cardiovascular disease ^7,8^. Even the quality of human aging has been related to mental health ^9,10^, but this association belies a mechanistic understanding of the relationship.

Despite its prevalence in our society, the mechanisms underlying death perception, the life-long consequences of such experiences, and the degree to which they are shared across species remain largely unknown. Reasons for this include an inherent difficulty conducting experiments on affected human populations and a dearth of animals models in which to identify causal relationships. In this study, we provide evidence that the vinegar fly, *Drosophila melanogaster*, has death perception; that this perceptive experience is driven by visual cues and modulated by olfactory cues; and that it induces both short- and long-term effects on health and longevity. Remarkably, the negative effects of death perception are reversed by targeted pharmacologic and genetic attenuation of serotonin signaling, suggesting deeply conserved links between psychological and physical health.

## Results

While investigating whether adult *Drosophila* behaviorally respond to diseased individuals in their environment, we discovered instead that flies become aversive after exposure to dead conspecifics. In our initial experiments, we established a binary choice assay (T-maze) in which flies that were previously infected with the lethal pathogen *Pseudomonas Aeruginosa PLCS* were placed behind a screen in one side of a T-maze and healthy flies were placed in the opposite side. When naïve choosers were loaded into the T-maze, we found that they sorted non-randomly, in that they avoided the side of the T-maze containing a group of flies that had been infected 24-48 hours previously (Supplementary Fig. 1a). We consistently failed to observe avoidance in naïve choosers to groups of flies that had been infected for less than 24 hours. Notably, flies began dying from our *Pseudomonas* infection roughly 24 hours post-infection, suggesting that the appearance of dead flies rather than infection *per se* might be the cause for the aversion. We therefore asked whether dead flies alone were sufficient to create an aversive stimulus. We found that they were not (Supplementary Fig. 1b). When comparing preference between only healthy live flies vs. only dead flies, naïve choosers preferred dead flies (Supplementary Fig. 1c) presumably due to CO_2_ emitted from live animals, which is a known repulsive stimulus^11^, establishing that the dead animals themselves were not intrinsically aversive. We therefore asked whether a mixture of dead flies with healthy live flies was aversive compared to healthy live flies alone. We observed a strong preference of naïve flies choosing the side of the T-maze without dead animals (Supplementary Fig. 1d). Finally, healthy flies from different laboratory strains that had been pre-exposed to dead conspecifics for 48 hours (the dead flies were removed immediately prior to the choice assay) exhibited aversive qualities (Fig. 1a and b), establishing that the presence of dead flies in the T-maze was not required for aversion. Together these data indicated that dead fly exposure triggered changes in healthy live individuals that repelled naïve choosers.

**Fig. 1.**
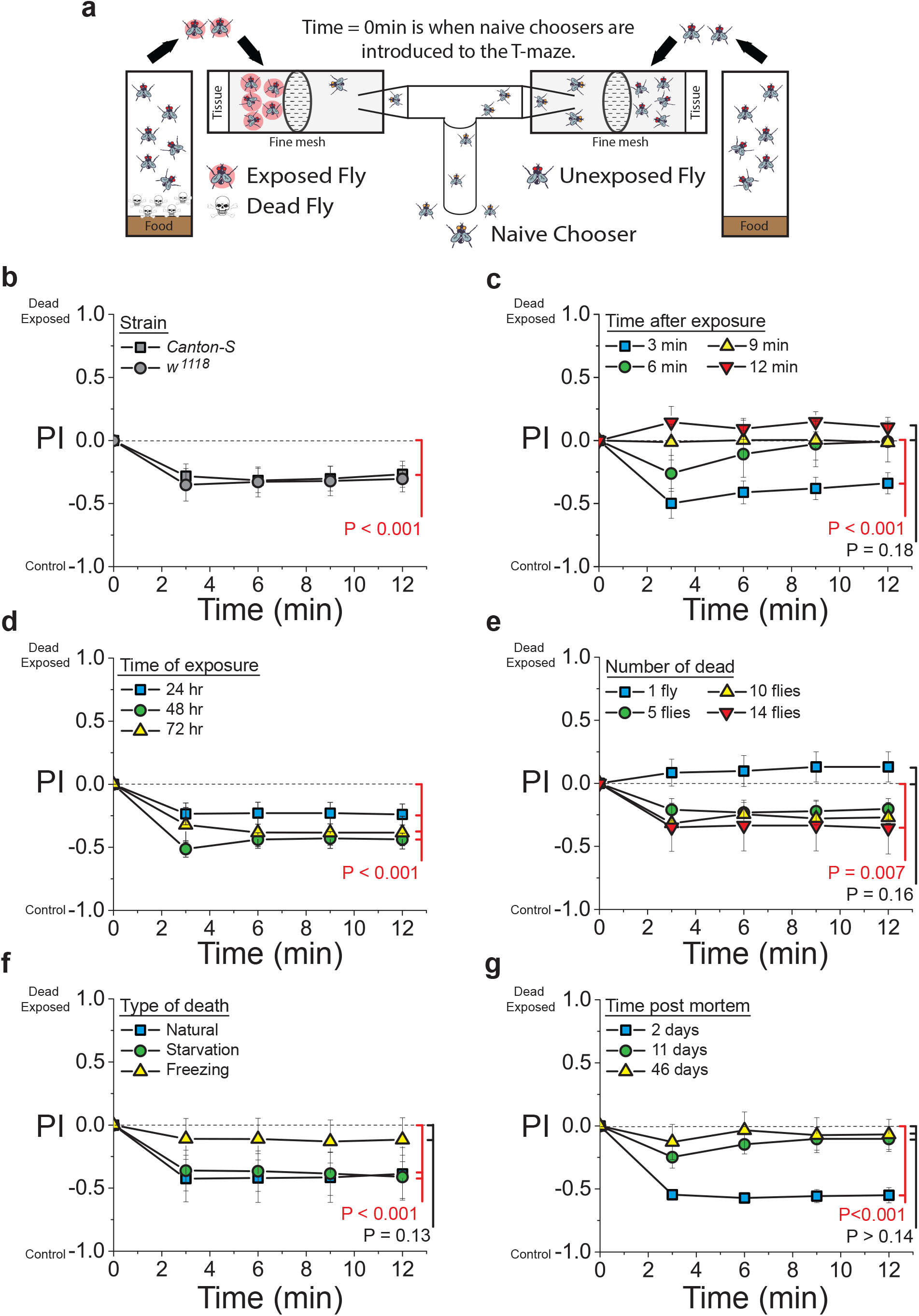
Flies become aversive after exposure to dead conspecifics. **a** Cartoon representing the exposure protocol and binary T-maze apparatus used in our choice behavior assays. PI = preference index calculated as (Number of flies in the exposed arm (N_E_)-Number of flies in the unexposed arm (N_C_))/total (N_C_+N_E_). **b** Flies of two different laboratory strains (Canton-S and *w^1118^*) that were exposed to dead conspecifics for 48 hours were aversive to naïve Canton-S choosing females (N = 9 for Canton-S and N = 7 for *w^1118^*, P < 0.001 for Canton-S and P = 0.001 for *w^1118^*). **c** Flies exposed to dead conspecifics retained their aversive characteristics to naïve choosing flies for up to 9 min after the dead flies were removed (N = 9 for each treatment, P = 0.01 for 6 min and P = 0.22 for 9 min, group ANOVA P < 0.001). **d, e** When flies were exposed to dead conspecifics, they evoked avoidance behavior in naïve choosing females that was intensified with (**d**) longer periods of exposure (N = 19 for 24 hours, N = 9 for 48 hours, and N = 14 for 72 hours, group ANOVA P = 0.027) and (**e**) the number of dead animals used during the exposure treatment (N = 6 for each treatment, P = 0.043 for 5 flies, P = 0.01 for 10 flies, group ANOVA P < 0.001). **f** Flies exposed to animals that died of natural or starvation-induced death, but not freezing death, evoked avoidance behaviors in naïve flies (N = 8 for natural and starvation-induced death and N = 10 for frozen induced death, group ANOVA P = 0.04). **g** Newly dead flies effectively induced the aversive cues in exposed animals, but long-dead flies did not (N = 6 for each treatment, P = 0.17 for 11 days dead and P = 0.14 for 46 days dead flies, group ANOVA P < 0.001). Except where noted in panel b, all naïve choosing female flies were from the Canton-S strain. Each T-maze sample tests 20 flies. Error bars represent SEM. All P-values were determined by non-parametric randomization (see Methods for details).

Many control experiments ruled out positional artifacts, biases in our technical apparatus, and bacterial infection as causes for the preferred segregation of naïve choosers away from flies exposed to dead animals. Naïve choosing flies segregated randomly when both sides of the T-maze were empty or when both sides contained equal numbers of live, unexposed flies (Supplementary Fig. 2a and b). Aversive cues were not induced when flies were mock-exposed to dead animals using black beads that are roughly the size of a fly, and the presence of dead flies did not affect feeding over 24 hours (Supplementary Fig. 2c and d). This suggests that aversive cues in exposed flies are not triggered by structured environments or by changes in food accessibility. We also observed no induction of aversive cues when the total number of flies in the pre-choice environments was equalized by augmenting the number of live flies in the no-exposure treatment, establishing that exposed flies are not exhibiting a response to perceived increases in population density (Supplementary Fig. 2e). Finally, similar levels of aversion were induced when axenic flies were used as both exposed and dead flies establishing that this effect was not caused by infection, bacterial proliferation, or changes in the gut biota in either dead or exposed animals (Supplementary Fig. 2f).

Subsequent experiments using only healthy animals pre-exposed to dead individuals before behavioral testing revealed that the effects of dead fly exposure are highly reproducible and are influenced by the characteristics of the dead individuals. Exposed flies lost their aversive characteristics approximately ten minutes after dead flies were removed (Fig. 1c), which is also roughly the span of short term memory in *Drosophila* ^12^, indicating that the aversive effect is persistent but short-lived. The aversiveness of exposed flies was also scalable by the number of dead: exposed flies became more aversive as the time of exposure and number of dead in the environment increased, although in standard rearing vials the magnitude of the effect saturated at roughly 48 hours and 10 dead animals, respectively (Fig. 1d and e). Flies exposed for 48hrs to flies that died from starvation or from normal aging triggered aversive cues, while a similar exposure to flies killed by immersion in liquid nitrogen did not (Fig. 1f). Long-dead flies also failed to induce aversive cues in exposed animals (Fig. 1g).

In humans, the severity of behavioral and emotional reactions to dead animals correlate positively with the extent that one identifies with the deceased ^13^. Insects often elicit little or no response, while mammals, especially those most like ourselves such as chimpanzees, can generate significant distress. Even among humans, physical or cultural similarities potentiate emotional responses ^14^. If the effects that we observed in flies shared aspects of our own death encounters, we would predict that the effects of such encounters would be influenced by the evolutionary relatedness between the dead and live animals. We tested this prediction by exposing *Drosophila melanogaster* to dead individuals from one of three related species *(Drosophila virilis, Drosophila simulans*, and *Drosophila erecta)*. We found that exposure to dead animals from the two closely related species *(D. simulans* and *D. erecta)* were able to induce aversive cues in *D. melanogaster* to a similar extent as did exposure to their conspecifics, while exposure to the evolutionarily more distant *D. virilis* did not (Fig. 2a).

**Fig. 2.**
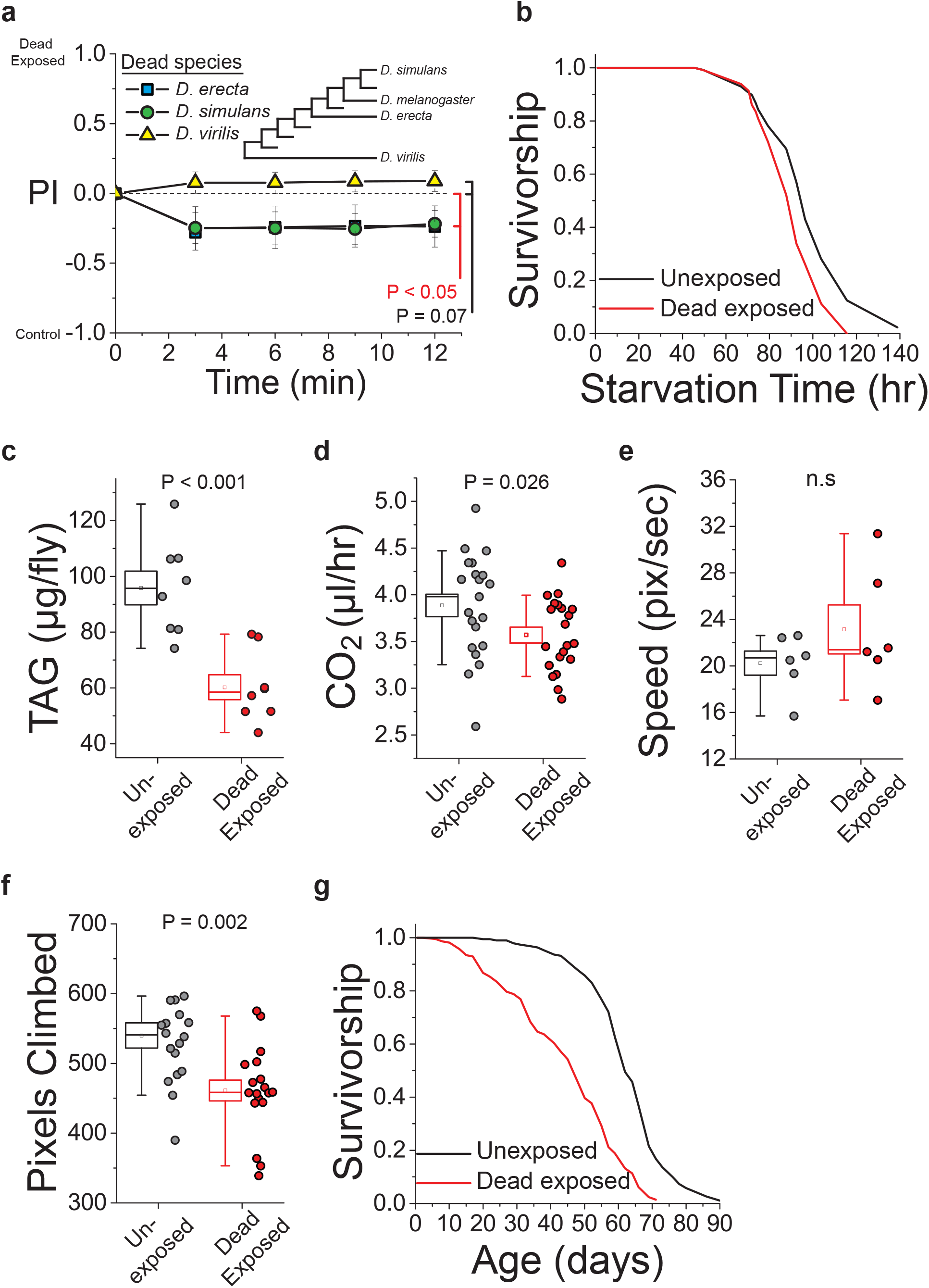
Exposure to dead conspecifics alters physiology and lifespan in *D. melanogaster*. **a** When *D. melanogaster* were exposed to dead animals from each of two closely related species *(D. simulans* and *D. erecta)* they presented aversive cues, but exposure to the evolutionarily more distant *D. virilis* had no effect (N = 10 for *D. erecta*, N = 8 for *D. simulans*, and N = 17 for *D. virilis*, P = 0.046 for *D. erecta* and P = 0.008 for *D. simulans*, group ANOVA P < 0.001). Inset depicts a phylogeny of related *Drosophila* species. **b, c, d** Flies exposed to dead conspecifics exhibited reduced (**b**) starvation survival (N = 100 per treatment, P < 0.001) (**c**) triacylglyceride stores (TAG, N = 8 biological replicates of 10 flies each), and (**d**) metabolic rate as measured by CO_2_ production relative to unexposed animals (N = 21 biological replicates of 5 flies/treatment). **e, f** While exposure to dead conspecifics did not affect (**e**) spontaneous movement rates, (N = 6 for each treatment, P = 0.26), (**f**) forced climbing was impaired relative to unexposed animals (N = 18 for each treatment). **g** Chronic exposure to dead animals significantly reduced lifespan flies (N = 190 for unexposed, 212 for exposed, P < 0.001). For panel a, naïve choosing flies were from the Canton-S strain. Each T-maze sample tests 20 flies. Error bars represent SEM. P-values for binary choice were determined by non-parametric randomization. Comparison of survival curves was via log-rank test, and the remaining phenotypes were evaluated for significance by t-test (see Methods for details).

Having observed that exposure of healthy flies to dead conspecifics consistently resulted in the production of aversive cues that repelled naïve choosers, we next sought to investigate whether this experience elicited broader physiological and health effects. We found that short-term exposure of *D. melanogaster* to dead conspecifics compromised starvation survival and reduced levels of triacylglycerol (TAG), which is the primary storage lipid in flies (Fig. 2b and c). It also resulted in a moderate but significant reduction in CO_2_ production, indicative of an altered metabolic rate (Fig. 2d). Exposed flies were capable of normal levels of spontaneous activity and exploration (Fig. 2e), but they showed impaired motivated climbing ability (Fig. 2f). Finally, chronic exposure to dead animals significantly reduced lifespan (Fig. 2g), which was robust to experimental strain (Supplementary Fig. 3a), was sex-specific in its magnitude (Supplementary Fig. 3b), was reduced in isolation (Supplementary Fig. 3c), and was not caused by population density (Supplementary Fig. 3d) or by environmental structure (Supplementary Fig. 3e).

We hypothesized that the effects of exposure to dead animals were driven by a perceptive experience that relied on one or more canonical sensory modalities in the healthy exposed flies. This is supported by the fact that gustatory and olfactory circuits have previously been shown to influence aging and physiology in *Drosophila* ^15-17^. We therefore asked which sensory modalities were necessary for aversivness to be triggered upon exposure to dead animals. We found that naïve choosers exhibited no behavioral preference in the T-maze when experimental animals were exposed to dead flies in the dark, indicating that this treatment failed to induce aversive cues (Fig. 3a). To rule out the unlikely possibility that light was somehow required to induce a potency in dead flies, we repeated these experiments in our standard 12h:12h light:dark conditions using *norpA* mutant flies, which are blind. We observed no evidence of aversive cues from *norpA* mutants following exposure (Fig. 3b). *Orco^2^* mutant flies are broadly anosmic, and they exhibited a significant, but not complete, loss of the aversive cues (Fig. 3c). On the other hand, flies lacking the *ionotropic receptor 76b*^18^, which is involved in chemosensory detection of amino acids and salt, exhibited normal aversion following dead exposure, as did flies lacking both *Ir8a* and *Ir25a* ionotropic co-receptors, which are required for multiple sensory functions ^19^ (Supplementary Fig. 5a-b). Flies carrying a *poxn* mutant allele, which are effectively taste-blind, exhibited a similar response to death exposure as genetically homogenous control flies (Fig. 3d).

**Fig. 3.**
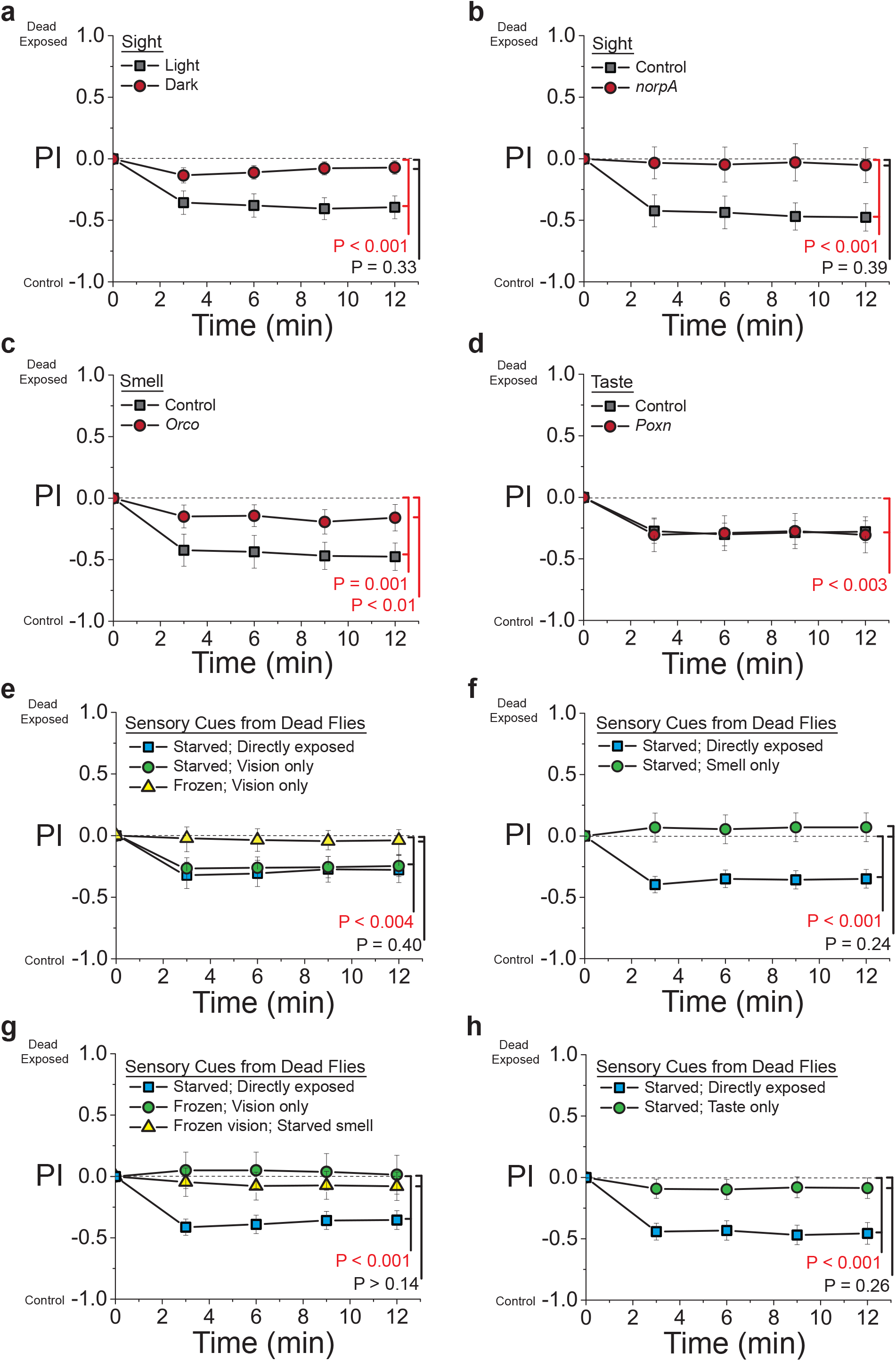
The avoidance behavior of exposure to dead conspecifics are mediated by sensory perception. **a** When experimental flies were exposed to dead animals in the dark, they failed to evoke avoidance behavior in naïve choosing females (N = 19 for light exposure and N = 20 for dark exposure). **b** Blind, *norpA* mutant flies maintained under light conditions also failed to exhibit the effects of death exposure on aversive cues (N = 8 for each treatment). **c** *Orco^2^* mutant flies, which are largely smell-blind, exhibited a near complete loss of aversive cues following exposure (N = 9 for *Orco^2^* and N = 8 for control). **d** Flies carrying the *Poxn^Δ4M22-B5-ΔXB^* mutation, which are effectively taste-blind, exhibited a similar effect of death exposure as control flies to aversive cues (N = 9 for each treatment, P = 0.002 for control and P < 0.001 for *poxn)*. **e** Canton-S mated female flies were exposed to flies that were dead either due to starvation or to liquid nitrogen-immersion for 2 days where the dead were kept beneath an acrylic floor on a thin layer of agar. This apparatus ensures that only visual cues are transferred. As a control treatment, a group of flies were directly exposed to starvation-induced dead for 48 hours in the absence of the acrylic barrier. Flies demonstrated avoidance behavior when exposed to starvation-killed dead but not liquid nitrogen-killed dead (N = 19 for starvation induced death, N = 20 for starved and frozen death vision only). **f** Live flies were co-housed with starvation-killed flies in the presence or absence of a mesh chamber that prevents physical interaction between the live and dead flies. Co-housing live flies with dead in the presence of the mesh chamber inhibited the avoidance behavior typically seen when dead-exposed flies are introduced to naive choosing females (N = 10 for each treatments). **g** Liquid nitrogen-induced dead flies that were placed on the top of the mesh chamber failed to evoke avoidance behavior in naïve-choosing females (Frozen; vision only). Placing starvation-induced dead below the mesh in addition to the liquid nitrogen-induced dead flies on top of the mesh also failed to cause avoidance behavior in naïve-choosing females (N = 8 for frozen only, N = 13 for starved and starved + frozen). **h** Starvation-induced dead flies were homogenized and spread on top of the food prior to exposure. Live Canton-S female flies were then exposed to either intact or grounded dead flies for 2 days. The ground-up dead flies failed to evoke avoidance behavior in naïve flies (N = 10 for ground up and control treatments). For binary choice assays, all naïve choosing flies were from the Canton-S strain. Each T-maze sample tests 20 flies. Error bars represent SEM. P-values for binary choice were determined by non-parametric randomization.

We next asked whether controlled exposure to different types of sensory cues from dead individuals was sufficient to induce aversiveness in healthy flies. Using a newly designed chamber that allowed experimental flies to see dead flies but remain physically separated from them (Supplementary Fig 6a), we found that the sight of starvation-killed flies was sufficient to induced aversive cues to the same extent as direct exposure (Fig. 3e). The sight of flies that had been killed by immersion in liquid nitrogen had no effect (Fig. 3e), thus replicating our earlier results (Fig. 1f) and indicating that flies have the visual acuity to distinguish differences in these two types of corpses, as humans do (Supplementary Fig 4a), and that flies killed by freezing lack one or more key visual characteristics that are present in flies that died naturally. The ability of flies to distinguish differences by sight may also explain why live *D. melanogaster* do not convey aversive signals when exposed to dead *D. virilis*, as these flies are darker and larger than *D. melanogaster* themselves (Supplemental Fig 4b). Interestingly, however, repeated exposure to flies killed by freezing over a 20 day (but not 10 day) period was sufficient to induce effects on aversiveness, indicating that flies may eventually learn to recognize these corpses or to respond to alternative cues (Supplementary Figs 6b-c). A second specialized chamber was used to investigate the sufficiency of olfactory cues (Supplementary Fig 6d) to induce aversion. No aversive cues were produced by flies that were exposed to volatile odors from naturally dead flies indicating that olfactory cues alone were not sufficient to induce averseness (Fig. 3f). Olfactory cues from naturally dead animals were also incapable of gating otherwise insufficient visual cues from animals that had died by freezing (Fig. 3g). Finally, experimental flies were unaffected by direct exposure to extracts from gently homogenized flies that had died by starvation, suggesting gustatory cues are also not sufficient to induce aversiveness (Fig. 3h).

Remarkably, the sensory requirements for exposure to dead conspecifics to influence aging are consistent with those that were observed to affect aversiveness. The significant reduction in lifespan that resulted from chronic exposure to dead animals was absent when flies were aged in constant darkness (Fig. 4a; Supplementary Fig. 7a). Notably, unexposed control flies were longer-lived in constant darkness, which is consistent with an effect of death perception in cohorts aging normally in light-dark cycles. Blind *norpA* mutants also exhibited significantly reduced effects on lifespan (Fig. 4b, Supplementary Fig. 7d). Long-term exposure to flies killed by freezing reduced lifespan (Supplementary Fig. 7b). Largely anosmic *Orco^2^* mutant flies exhibited a partial decrease of the effect on lifespan (Fig. 4c), as did flies with their antennae removed (Supplementary Fig. 7c), while flies lacking *Ir76b* or flies lacking both *Ir8a* and *Ir25a* exhibited a decreased lifespan following dead exposure to the same extent as genetically homogenous control animals (Supplementary Fig. 5c-d). Finally, *poxn* mutants also exhibited a similar decrease in lifespan in response to death exposure as control flies (Fig. 4d, Supplementary Fig. 7d).

**Fig. 4.**
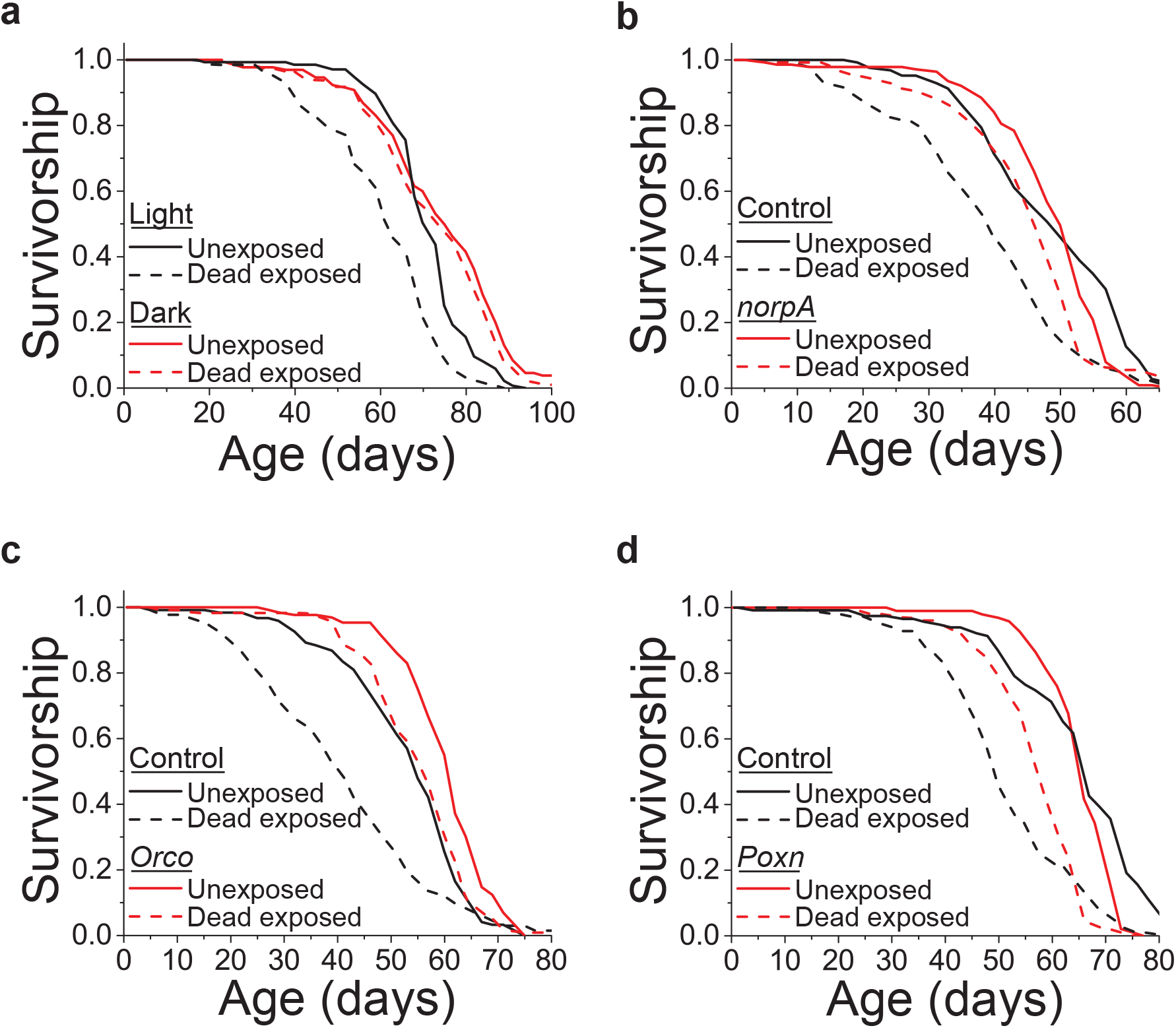
Exposure of dead conspecifics causes changes in lifespan that are mediated by sight and smell. **a**, Exposure of live animals to dead in the light, but not the dark, affected lifespan (N = 118 for dark dead exposed, 130 for dark unexposed, 125 for light dead exposed, and 135 for light unexposed, P < 0.001 for light and P = 0.18 for dark, P < 0.001 for the interaction between light and exposure via Cox Regression). **b**, The lifespan effect observed in *norpA* mutant flies following exposure to dead conspecifics was significantly diminished relative to control animals (N = 139 for *norpA* dead exposed, 126 for *norpA* unexposed, 126 for control dead exposed, and 131 for control unexposed, P < 0.001 for control and P = 0.003 for *norpA*, P < 0.001 for the interaction between genotype and exposure via Cox Regression). **c**, Anosmic flies maintained a reduced effect of death exposure on lifespan compared to control flies (N = 115 for *Orco^2^* dead exposed, 129 for *Orco^2^* unexposed, 133 for control dead exposed, and 121 for control unexposed, P < 0.001 for control and *Orco^2^*, P = 0.002 for the interaction between genotype and exposure via ANOVA). **d**, Taste-blind flies showed normal lifespan effects due to dead exposure (N = 101 or *Poxn* dead exposed, 96 for *Poxn* unexposed, 112 for control dead exposed, and 116 for control unexposed, P < 0.001 for control and *Poxn*, P = 0.06 for the interaction between genotype and exposure via Cox Regression). Survival curve comparison was accomplished using a log-rank test (see Methods for details).

Together, these data indicate that there is an essential perceptual component associated with the physiological and health effects of exposure to dead conspecifics. Sight of naturally dead conspecifics is both necessary and sufficient to induce physiological effects, suggesting a model in which visual cues serve as the primary way in which *D. melanogaster* distinguish their dead and adding to a growing literature indicating that flies are capable of extracting and responding to different ecologically-relevant visual cues in their environment, such as parasites and competitors ^20,21^. While gustatory cues are apparently not involved in the effects that we observe, the role of olfaction is less clear. Smell-deficient flies respond less strongly to dead individuals, but odors from dead flies are not sufficient to induce measurable changes in aversiveness. Interestingly, the aversive cues emitted by healthy flies following exposure to dead individuals have a significant olfactory component: when *Orco^2^* mutants were used as naïve choosers in the T-maze assay (e.g., Fig. 1), they assorted randomly between the two arms (Supplementary Fig. 8). Similar results were observed when naïve choosers carried a loss of function mutation for *Gr63a*, an essential component of the *Drosophila* CO_2_ receptor (Supplementary Fig. 8). We therefore currently favor a model in which olfaction mediates a social cue among exposed animals that magnifies the effects of visual death perception.

We next asked whether we might identify a neural signature of this putative perceptive experience by comparing the neuro-metabolome of experimental flies following 48 hours of dead exposure to that of unexposed animals. *norpA* mutant flies, which were treated identically but which lacked the perceptive experience, were analyzed simultaneously to account for temporal effects and to isolate potential causal metabolites. Targeted metabolite analyses identified 119 metabolites present in treatments for positive and negative modes (see Supplementary file for raw data). Using a randomization procedure together with principal component analysis (PCA), we identified a single principal component (PC10) that significantly distinguished the neuro-metabolomes of exposed and unexposed flies but was unchanged by exposure in *norpA* mutant flies (Fig. 5a and Supplementary Fig. 9a). Only one metabolite (glyceraldehyde) was found to be statistically correlated with exposure to dead individuals in experimental but not *norpA* mutant flies (Fig. 5b). This low number is likely due to limited statistical power to reveal significant interactions. However, the multivariate analysis revealed an intriguing trend: five of the top ten metabolites associated with the effects of death perception (i.e., those strongly loaded on PC10) have been implicated in anxiety, depression, and/or mood disorders in humans (specifically lactate ^22^, quinolinate ^23^, sorbitol ^24^, 3-hydroxybutyric acid ^4^, and sarcosine ^9^).

**Fig. 5.**
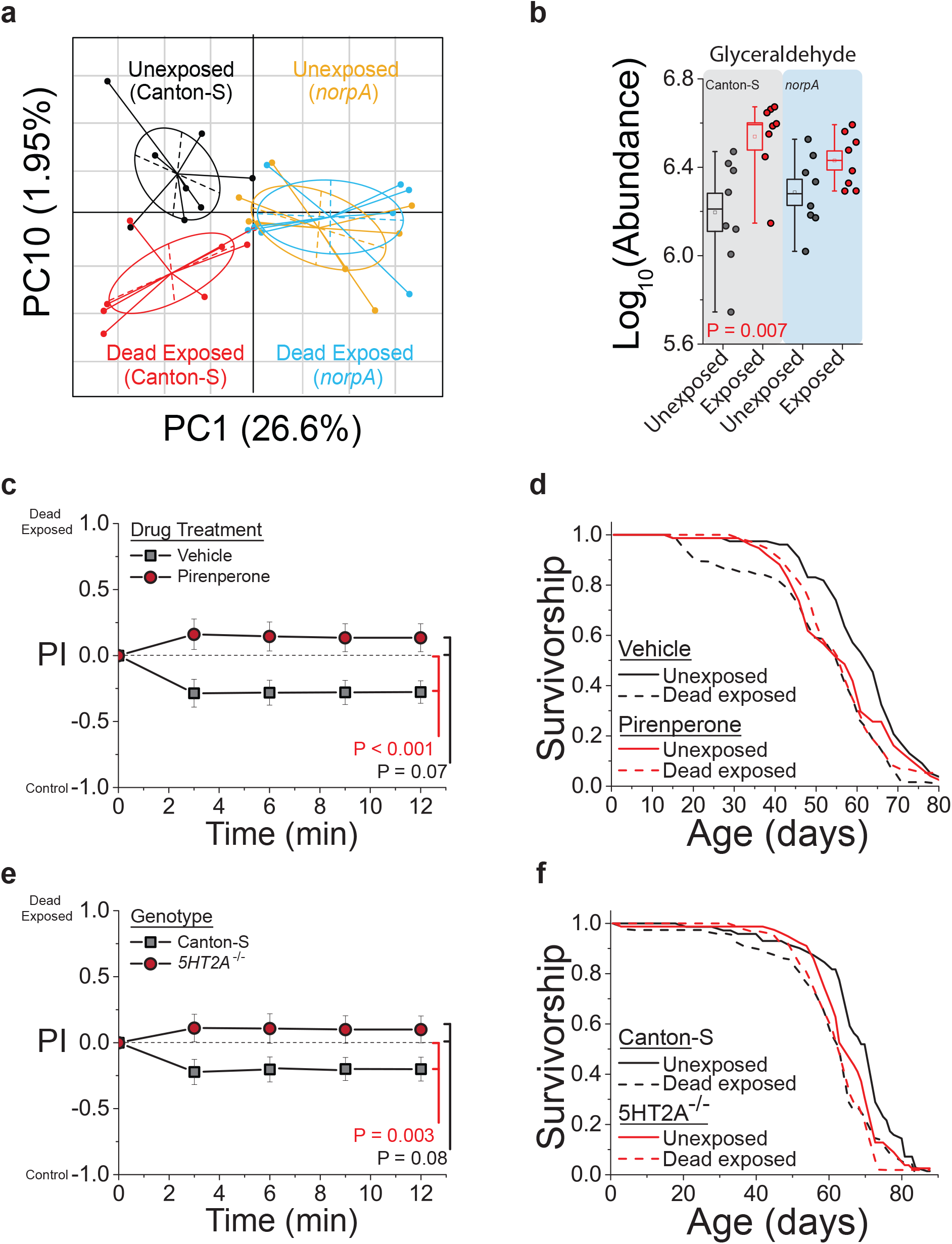
Death perception elicits acute changes in the neuro-metabolome, and its effects on health are mitigated by manipulations that attenuate serotonin signaling. **a** Principal component plot showing the distribution of samples for each treatment. Neuro-metabolites weighted heavily in PC10 distinguish the effect of death exposure, while those favored in PC1 distinguish genotype. Plots represent mass spectrometry analysis of metabolites identified under positive mode (N = 8 biological replicates, with 40 fly heads per replicate). **b** Glyceraldehyde abundance was significantly increased in flies following death perception, but it was unchanged in blind *norpA* mutant flies similarly treated. **c, d** Pharmacologic treatment of Canton-S females with the serotonin receptor 5HT2-antagonist, pirenperone, during exposure to dead conspecifics effectively protected them from the consequences of death perception on (**c**) aversive cues detected by naïve choosing flies (N = 10 for each treatment) and (**d**) lifespan (N = 75 for pirenperone-fed dead exposed, 76 for pirenperone-fed unexposed, 75 for vehicle-fed dead exposed, and 77 for vehicle-fed unexposed, P < 0.001 for vehicle-fed and P= 0.86 for pirenperone-fed, P = 0.007 for the interaction between drug and exposure via ANOVA). **e, f** Null mutation of serotonin receptor 5-HT2A protected flies from the consequences of death perception on (**e**) aversive cues detected by naïve choosing flies (N = 15 for each treatment) and (**f**) lifespan (N = 63 for *5-HT2A^-/-^* dead exposed, 78 for *5-HT2A^-/-^* unexposed, 75 for Canton-S dead exposed, and 72 for Canton-S unexposed, P < 0.001 for Canton-S control flies and P= 0.06 for *5-HT2A^-/-^* mutants). A replicate lifespan experiment revealed the same results (see Supplemental Fig. 5c), and P = 0.05 for the combined interaction between genotype and exposure via ANOVA. P-values for principal component analysis and for binary choice were determined by non-parametric randomization. Each T-maze sample tests 20 flies. Error bars represent SEM. Comparison of survival curves was via log-rank test, and individual metabolites were evaluated for significance by t-test (see Methods for details).

Following the neuro-metabolomic analysis, we examined several candidate pharmacologic compounds with the goal of identifying molecular targets that are required for the health consequences of death perception. We chose a small number of compounds that were related to those used for treatment of human patients suffering from anxiety, depression, or the consequences of severe traumatic events. For T-maze assays, flies were treated with each drug prior to exposure to dead individuals, while for lifespan assays flies were treated throughout life. Of those compounds examined, we found one, pirenperone, which abrogated the effects of death perception on both aversive cues and lifespan (Fig. 5c and d, Supplementary Table 1). Pirenperone is a putative antagonist of the serotonin 5-HT2 receptor in mammals, although it may interact with other biogenic amine receptors at high concentrations ^25^. We therefore tested whether loss of the 5-HT2A receptor recapitulated the effects of pirenperone and abrogated the effects of death perception in *Drosophila*. We found that it did (Fig. 5e and f, Supplementary Fig. 9b and c), suggesting that serotonin signaling through the 5-HT2A branch is required to modulate health and lifespan in response to this perceptive experience.

Finally, we asked whether activation of *5-HT2A+* neurons was sufficient to recapitulate the aversion and lifespan phenotypes we observed following death perception. We ectopically expressed the thermosensitive cation channel Transient Receptor Potential A1 (TRPA1) in cells that putatively produce 5-HT2A (using *5-HT2A-GAL4)*. The *Drosophila* TrpA1 channel promotes neuron depolarization only at elevated temperatures (>25°C), thereby allowing temporal control over cell activation ^26^. Experimental flies *(5-HT2A-GAL4>UAS-dTrpA1)* and their genetic control strain *(5-HT2A-GAL4;+)* were grown and maintained for 10 days after eclosion at 18°C (non-activating conditions). They were then placed at 29°C for 48 hours to mimic death exposure. We found that after activation, experimental flies were significantly more aversive relative to the genetic control (Fig. 6a; P < 0.001). We were also able to mimic the effects of chronic death exposure by activating *5HT2A*^+^ neurons throughout life, when we observed a rapid and significant increase in death rates consistent with our previous data (Fig. 6b).

**Fig. 6.**
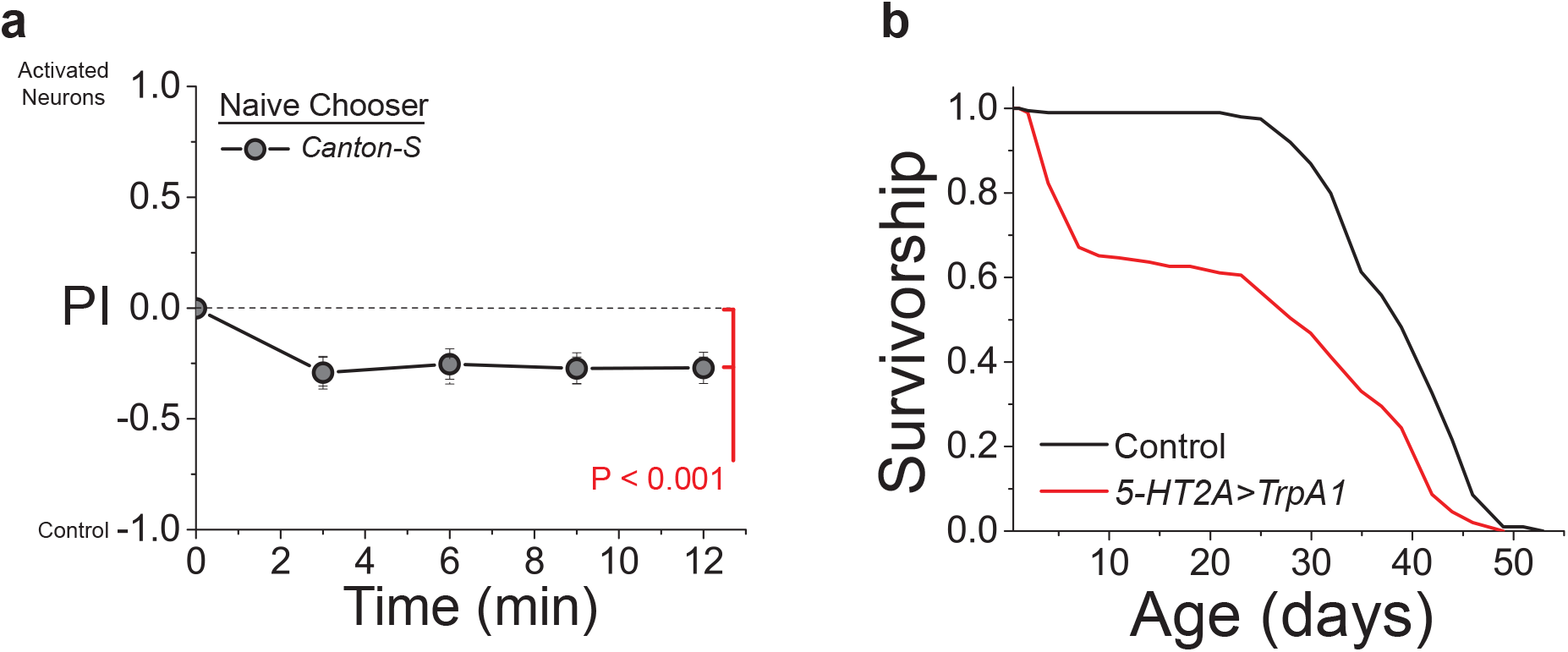
Activation of 5-HT2A neurons induces aversiveness and reduces lifespan in *Drosophila*. Constitutive activation of *5-HT2A^+^* neurons via expression of *UAS-TrpA1* relative to *5-HT2A-GAL4;+* control flies results in (**a**) increased aversiveness (N = 19, P < 0.001) and (**b**) significantly decreased lifespan (N = 194 for *5-HT2A>TrpA1* and N = 195 for *5-HT2A-GAL4;+*, P < 0.001). Each T-maze sample tests 20 flies. Error bars represent SEM. Comparison of survival curves was via log-rank test.

## Discussion

We believe that these data provide evidence that, for *Drosophila melanogaster*, the perception of dead conspecifics in the environment creates a psychological experience that is transduced by conserved signaling pathways into behavioral and physiological changes that influence health. We speculate that dead conspecifics serve as a threat cue and that long-term exposure leads to a form of mental stress that has negative consequences on metabolism, physical condition, and aging ^27,28^. Associations linking human morbidity and disease with dramatic psychological events and subjective assessments of “well-being” are prevalent in the demographic and epidemiological literatures ^29^, and our results may provide insight into their underlying molecular mechanisms.

Serotonin is one of the most well-studied and influential neurotransmitters in multiple organisms, including humans, and this report adds to growing evidence that it is an important component of how different sensory experiences modulate aging and aging-related disease via specific neural signaling pathways across taxa. In *C. elegans*, a global, cell non-autonomous response to heat is triggered by thermo-sensory neurons in a serotonin dependent manner, which is also capable of extending lifespan ^30^. Lifespan extension by activation of the hypoxic response also requires specific components of serotonin signaling in sensory neurons ^31^. In *Drosophila*, serotonin is required for protein perception, and loss of receptor *5-HT2A* increases fly lifespan in complex and potentially stressful nutritional environments ^32^. Whether serotonin is merely permissive for changes that affect lifespan or whether it directly modulates aging remains unclear. In *C. elegans* there is evidence for a direct role, as feeding of serotonin receptor antagonists is sufficient to extend lifespan by putatively mimicking dietary restriction^33^. We also present evidence that activation of 5-HT2A neurons is sufficient to modulate aging, but these results should be interpreted with caution; both aversiveness and short lifespan might instead be reflective of unrelated molecular changes that result in less healthy animals.

While death perception is known to occur in some species throughout the animal kingdom ^4^, this is, to our knowledge, the first indication that such an ability may be present in an invertebrate laboratory model system. *Drosophila* neuroscience and genetics may now be directed toward unraveling how flies recognize dead animals and how this specific perceptive experience triggers physiological changes in peripheral tissues. Work in model systems has pioneered interest in how the brain uses sensory information to control aging through systemic changes in stress resistance and nutrient utilization ^34-36^. These same systems may also allow us to probe the mechanisms underlying how aging and other health metrics are determined by fundamental brain processes that are historically rooted in psychology, such as perception and cognition, and that have long been considered uniquely human ^37^

## Methods

### Contact for Reagent and Resource Sharing

*Further information and requests for resources and reagents should be directed to and will be fulfilled by the Lead Contact, Scott Pletcher*.

### Experimental Model and Subject Details

The laboratory stocks *w^1118^*, Canton-S, *UAS-dTrpA1, norpA*, and *Ir76b^-/-^* [BL51309] *Drosophila* lines were obtained from the Bloomington Stock Center. *Poxn^ΔM22-B5ΔXB^* and *Poxn^Full1^* were provided by J. Alcedo ^38^. *Orco^2^* mutant flies were a generous gift from L. Vosshall ^39^. *Gr63a^1^* mutant flies were a gift from A. Ray. *5-HT2A^PL00052^* mutant and *5-HT2A-GAL4 (3299-GAL4)* flies were graciously provided by H. Dierick. *Ir8a^-/-^ /Ir25a^-/-^* mutant flies were kindly provided by R. Benton. Three species of *Drosophila (D. simulans, D. erecta* and *D. virilis)* were generously provided by P. Wittkopp. All of these strains were maintained on standard food at 25°C and 60% relative humidity in a 12:12 h light:dark cycle.

### Method Details

#### Generation of dead flies

Unless otherwise noted, dead flies were generated by starvation. One-to two-week old Canton-S female flies were separated using CO_2_ anesthesia and transferred to vials containing 2% agar. Flies were transferred to fresh agar vials every ~3 days and dead flies were collected within three days of death. Vials in which dead flies stuck to the agar were not used. For the data in Fig 1F, flies died from natural causes or were killed by rapid freezing in liquid nitrogen. Age-matched Canton-S females were used for rapid freezing.

#### Short-term exposure to dead flies

Twenty 2-week-old mated female flies were collected under light CO_2_ anesthesia and exposed for 48 hours to 14 freshly dead female flies in standard food, where they freely interacted with the dead flies. During the 48 hour exposure period, flies were maintained in a 12:12 hour light:dark cycle, except for those involving exposure in 100% darkness, which took place in a closed incubator. In both cases, flies were maintained at 25°C and 60% relative humidity. To test species specificity, experimental female flies (*D. melanogaster*) were exposed to three different species of dead *Drosophila*; we used 14 freshly dead female flies of *D. melanogaster, D. simulans*, and *D. erecta*, and 8 freshly dead *D virilis* due to their larger size.

#### Behavioral preference assays

To generate naïve choosers for preference assays, newly eclosed virgin female flies (< 7 hours old) were collected and transferred to standard food vials with 3 male flies per 20 females. Flies were kept at 25°C and 60% humidity, with a 12:12 hour light: dark cycle for 2 days, after which they were briefly anaesthetized to remove the male flies. Mated female flies were then transferred to fresh vials for one day to recover. On day four post-eclosion, the flies were placed into vials containing moist tissue paper for 4 hours prior to their introduction into the T-maze for behavioral monitoring. Choice was measured using binary traps made from commercially available T connectors (McMaster-Carr Part Number <u>5372K615</u>) with 200μl pipette tips, which were trimmed, attached to opposite ends of the T connectors to form one-way doors that end in small collection chambers. Experimental flies that were pre-exposed to dead flies were loaded into a collection chamber with moist tissue paper in one arm of the T-connector while unexposed flies were loaded into the opposite chamber that also contained moist tissue paper. Unless otherwise noted, 20 live flies were exposed to 14 dead animals. Twenty naïve choosing flies were introduced into the central arm of the maze, and the number of flies trapped in each arm were counted at regular intervals. Behavioral preference was measured in a dark room under dim 660 nm red light at 24°C, and behavior was observed at 3, 6, 9, and 12 mins. A Preference Index (PI) at each time point was computed as follows: (Number of flies in exposed arm (NE)-Number of flies in unexposed arms (N_C_))/(N_C_+N_E_). The fraction of flies that participated in the experiments was calculated as: (N_C_ + N_E_)/20. Average PI values are weighted mean values among replicates with weights proportional to the number of animals that made a choice. Participation rates for all of the T-maze assays were > 50%. Experiments were replicated at least 2 independent times. Beads used for mock dead flies were obtained from Cospheric innovation in Microtechnology (Catalog number: CAS-BK 1.5mm).

#### Starvation experiments

At day 4 post-eclosion, 10 mated female flies were separated in 10 vials/treatment containing 2% agar. Flies were kept in constant temperature and humidity conditions with a 12:12 hour light: dark cycle. A census of live flies was taken every 6-8 hours. For exposed flies, the dead flies were left in each vial throughout the experiment. For control flies, dead flies were removed at each census point. Flies were transferred to fresh agar vials every 6 hours.

#### TAG assays

Four-day old, adult Canton-S female flies were collected and subsequently handled using our standard short-term exposure protocol (see above). Following the 48 hour exposure to dead animals, live experimental flies were removed and homogenized in groups of 10 in 150 μl PBS/0.05% Triton X. Unexposed flies were collected simultaneously. The amount of TAG in each sample was measured using the Infinity Triglyceride reagent (Thermo Electron Corp.) according to the manufacturer’s instructions. Eight independent biological replicates (of 10 flies each) were obtained for treatment and control cohorts.

#### Negative geotaxis assay

Four-day old, adult Canton-S female flies were collected and subsequently subjected to our standard short-term exposure protocol (see above). Following the 48 hour exposure to dead animals, live experimental flies and their corresponding unexposed controls were removed and transferred to climbing chambers by aspiration. Negative geotaxis was measured using DDrop, an automated machine developed in the Pletcher laboratory that drops flies from 24” and then tracks upward movement of individual flies through a video tracking algorithm. For each fly, we calculated both the total distance travelled and the time required to reach individual quadrants of the chamber.

#### CO_2_ measurement

Four-day old, adult Canton-S female flies were collected and subsequently subjected to our standard short-term exposure protocol (see above). Following the 48 hour exposure to dead animals, live experimental flies and their corresponding unexposed controls were removed, and CO_2_ production was measured from groups of five female experimental flies alongside their corresponding unexposed controls at 25°C. We used a Sable Systems Respirometry System, including a LiCor LI-7000 carbon dioxide analyzer, a Mass Flow Controllers (MFC2), and a UI-2 analog signal unit. Immediately prior to analysis, flies were transferred without anesthesia into glass, cylindrical respirometry chambers. Flies were allowed to acclimate to the new environment for 8 min before CO_2_ collection began. Six chambers were analyzed simultaneously using stop-flow analysis and the Sable Systems multiplexer. Incoming atmospheric air flow was dried, scrubbed of CO_2_, and then rehydrated before entering the respirometry chambers via the multiplexer. For each group, we collected 3 measures of CO_2_ production over a period of 20 min that were averaged to determine a final, single estimate of CO_2_ production per group. CO_2_ production values were obtained using the EXPDATA software from Sable Systems, following adjustment using a proportional baseline.

#### Video tracking

Mated Canton-S female flies were starved for 4 hours in a vial with moist tissue paper prior to the experiment to encourage movement within the chambers. Each chamber contained two dumbbell-shaped arenas comprised of two circles (Internal diameter = 1.0”) separated by a narrow corridor connecting them. A thin 2% layer of agar served as the floor of the chambers. Dead flies were lightly pressed into the agar of 1 arena to secure them. The positions of these stimuli were randomized within each experiment. Five minutes prior to recording, single exposed flies were loaded into the chambers by aspiration. Movement in each arena was recorded for 2 hours in a 25°C incubator under white light. Recordings were analyzed using the DTrack Software, developed in the Pletcher laboratory ^40,41^. From the tracking data, we calculated the amount of time each fly spent in each side of the arena, which was then used to calculate the relevant Preference Index (PI).

#### Feeding analysis

Feeding behavior was measured using the Fly Liquid Interaction Counter (FLIC) as described previously ^32^. Following our standard 48 hour exposure treatment, individual Canton-S female flies were placed into a single FLIC chamber with two food wells, each containing a 10% sucrose solution. Two independent experimental blocks were conducted using 15 dead exposed and 15 unexposed flies per experiment, providing a total of 30 flies per treatment. The experiments were performed at constant temperature (25°C) with 12:12 hour light: dark cycle. Throughout the experiment, three dead flies were kept in the chambers assigned to the dead exposed condition. Feeding interactions with the food were measured for 24 hours continuously using the FLIC reservoir system (see http://www.wikiflic.com). Data were analyzed using the FLIC Analysis R Source Code (available from wikiflic.com). Relevant feeding measures included the number of total interactions with the food, the total time spent interacting with the food, mean duration of each putative feeding event, and mean time between feeding events. These data were determined to be normally distributed, and a t-test was used to determine whether statistically significant differences were observed after noting the absence of significant block effects.

#### Antennal dissection

For experiments in which flies had the antenna removed, two-day old mated CS female flies were lightly anesthetized using CO_2_. Antennae were removed by fine forceps (Tweezers Dumont #55, WPI), after which the flies were transferred to bottles containing 10% sugar/yeast (SY) food. Antennae-less flies were held in an incubator at 25 °C with a 12:12 hour light dark cycle and 60% relative humidity for four days for surgical recovery. On day 7 post-eclosion flies were transferred to fresh SY10 vials for lifespan experiments.

#### Axenic Fly Culturing

Canton S flies were placed into cages with purple grape agar and yeast paste for approximately 18 hrs. Embryos were then collected with 10ml PBS and moved into a sterile hood, where they were treated with 10ml 1:10 sterile bleach solution (3x washes) and then washed with sterile water. Using sterile technique, 8ul of embryos were aliquoted into sterile 50ml falcon tubes that contained 8ml sterile standard fly media. Embryos were allowed to develop in a humidified incubator at 25°C with 12hr:12hr light:dark cycles. After 12 days, axenic flies were collected in a sterile hood into fresh, sterile 50ml falcon tubes containing 8ml sterile standard fly media. Flies were aged for 2 weeks, during which time fresh, sterile media was provided every 2-3 days. After 2 weeks, the flies were split into 2 groups with half of the group given fresh, sterile agar (to generate sterile dead flies) and the other half given fresh, sterile standard fly media. Sterile dead exposures occurred in the sterile hood using sterile tools and sterile technique. Control, conventionally-reared flies (non-sterile) were handled in an identical manner, with the exception of bleach washing. The sterility of the axenic flies was verified by plating the supernatant from total fly extracts onto brain heart infusion agar plates; colonies grew from the extracts of traditionally-reared flies but never from the extracts of sterile flies.

#### Neurometabolomic analysis

Following 48 hours of dead fly exposure, experimental flies were quickly frozen in a dry ice bath, and stored at −80°C overnight. Heads were removed via vortexing and manually separated from the body parts. Forty heads were then homogenized for 20 sec in 200μl of a 1:4 (v:v) water:MeOH solvent mixture using the Fast Prep 24 (MP Biomedicals). Following the addition of 800 μl of methanol, the samples were incubated for 30 mins on dry ice, then homogenized again. The mixture was spun at 13000 RPM for 5 mins at 4°C, and the soluble extract was collected into vials. This extract was then dried in a speedvac at 30°C for approximately 3 hours. Using a LC-QQQ-MS machine in the MRM mode, we targeted ~200 metabolites in 25 important metabolic pathways, in both positive and negative MS modes. After removing any metabolites missing from more than 5 out of 32 samples (15%), we were left with 119 metabolites. Metabolite abundance for remaining missing values in this data set were log-transformed and imputed using the k-Nearest Neighbor (KNN) algorithm with the impute package of R Bioconductor (www.bioconductor.org). We then normalized the data to the standard normal distribution (μ=0, σ^2^=1). Principal Component Analysis (PCA) was performed using the made4 package of R Bioconductor. We used permutation tests (n=10,000) to select PCs that significantly separate between different treatments (genotype and/or exposure to dead flies). For each permutation, we randomly distributed the treatments to the real abundance of each metabolite. PC analysis was done for both randomized and real data. The degree of separation for each PC can be measured by analyzing between- and within-group variance based on the projection of samples on that PC, which is indicated by the Z-score:

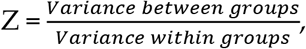

Variance between groups 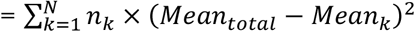, where N indicate the number of groups and *n_k_* indicates the number of samples in Group k. The distribution of Z-score was obtained from 10,000 randomized datasets. PCs that significantly deviated from this randomized distribution were considered as a significant separation of groups.

To identify individual metabolites of interest that are likely to be associated with death exposure, we sorted them per loadings on PC10 and selected the Top 10. For these candidates, we looked for metabolites that were (*i*) significantly different between control flies exposed to dead conspecifics versus those not exposed and that were (*ii*) less affected in *norpA* mutant flies. We found glyceraldehyde significantly up-regulated upon death exposure in Canton-S flies (one-sided Student’s t-test, P = 0.007), whereas such differences were not significant in *norpA* flies (one-sided Student’s t-test, P = 0.07).

#### Survival experiments

For lifespan experiments, experimental and control flies were reared under controlled larval density and collected as adults within 24 hours of emergence onto standard food where they were allowed to mate freely for 2-3 days. At 3 days post-eclosion, female flies were sorted under light CO_2_ and placed into fresh food vials. Contrary to our short-term protocol, experimental flies were chronically exposed to dead animals throughout their life. Lifespan measures were obtained using well-established protocols ^42^. Flies were transferred to fresh food vials every Monday, Wednesday, and Friday, at which time 14 freshly dead flies (~2-3 days old) were added. Unexposed animals were transferred simultaneously, but instead of adding dead flies we removed any flies that had died since the last census time. Flies were maintained at 25°C and 60% humidity under a 12:12 hour light: dark cycle. For experiments in the dark, flies were maintained in a dark incubator at 25°C and 50% humidity. Vials were changed as described above, and dead flies were counted under dim red light.

#### Drug administration

All drugs were purchased from Sigma-Aldrich. Each drug was initially dissolved in 100% DMSO at 10 mM concentration, aliquoted, and stored at −20°C. Every Monday, Wednesday, and Friday, an aliquot of the drug stock was thawed and diluted 1:500 in water for lifespans (20 μM final concentration). A similar dilution of DMSO alone was made in water as a vehicle control. Then, 100 μl of the diluted drug or vehicle control was added to each vial, coating the top of the food surface. After the liquid evaporated (~2 hours), the vials were ready for use as described above. For the behavior assay, flies were pretreated with 1 mM pirenperone or an equivalent dilution of DMSO for 2 weeks prior to exposure, in the absence of dead flies.

#### 5-HT2A Activation Methods

*5-HT2A-GAL4 (3299-GAL4*, H. Dierick, BCM) flies were crossed to *UAS-dTrpA1* to generate *5-HT2A>dTrpA1* flies. *UAS-dTrpA1* was backcrossed for at least 10 generations to *w^1118^* and therefore *5-HT2A-GAL4;+* was used as a genetic control strain. For behavioral assays, progeny from all crosses were maintained at 18°C until they were 10-14 days old, after which they transferred to 29°C for 48 hours prior to mimic exposure to dead individuals. The T-maze assay was established with *5-HT2A>dTrpA1* on one side of the assay and *5-HT2A-GAL4;+* on the other. Canton-S flies were used as naïve choosers. For lifespan experiments, progeny from both crosses were maintained at 18°C throughout development. Following eclosion, females were mated for 3 days, separated by gender, and placed at 29°C to begin lifespan measurement.

#### Bacterial infection with P. aeruginosa

The PA14 *plcs* strain used in this study was obtained from L. Rahme (Harvard Medical School). For each experiment, a glycerol stock was freshly streaked onto an LB/gentamycin plate. After an overnight incubation, a single colony was picked and grown in 1ml of LB/gentamycin until this seed culture reached logarithmic phase. Subsequently, the culture was diluted in 25 ml of LB/gentamycin and grown until the desired A_600_ concentration was reached. Finally, the bacterial culture was centrifuged and the pellet resuspended in LB media to obtain an A_600_ reading of 100. The culture was kept on ice during infection. Needles were directly placed in the concentrated bacterial solution and then poked into the fly abdomen. After infection, flies were transferred to standard food vials and kept in the incubator at 25°C and 60% humidity. Flies were collected 24 hours post infection for behavioral experiments. Infected flies were loaded in one arm of the T-connectors and control flies (not infected with *P. aeruginosa*) were loaded into the opposite arm. Twenty naïve choosing flies were introduced into the central arm of the maze and the number of flies in each arm of the trap was counted at regular intervals.

### Quantification and Statistical Analysis

For all preference assays, P-values comparing the Preference Index among treatments was obtained using a randomization procedure and the statistical software R. Briefly, the null distribution of no difference among treatments was obtained by randomizing individual preference indices obtained from groups of 20 flies among all measures (maintaining block structure when appropriate) and 100,000 t-statistics (or F statistics for multiple comparisons). P-values (one-sided or two-sided as appropriate) were determined by computing the fraction of null values that were equal or more extreme to the observed t-statistic (or F-statistic). Mean preference values were plotted and weighted by the number of choosing flies in each trial, with the error bars representing the standard error of the mean. Experiment-wise error rates for experiments comparing three or more treatments were protected by presentation of treatment P-value from non-parametric, randomization ANOVA, which are reported in the Figure Legends when appropriate. For lifespan and starvation assays, we employed survival analysis. Unless otherwise indicated, group- and pairwise-comparisons among survivorship curves (both lifespan and starvation) were performed using the DLife computer software ^42^} and the statistical software R. P-values were obtained using log-rank analysis (select pairwise comparisons and group comparisons or interaction studies) as noted. Interaction P-values were calculated using Cox-Regression when the survival data satisfied the assumption of proportional hazards. In other cases (as noted in the figure legends), we used ANOVA to calculate P-values for the interaction term for age at death. For all box plots, the box represents Standard Error of the Mean (SEM, centered on the mean), and whiskers represent 10%/90%. For CO_2_, TAG, and negative geotaxis measures, P-values were obtained by standard two-sided t-test after verifying normality and equality of variances. Details of the metabolomics analysis are presented above.

### Data Availability

Metabolomics data and analyses are provided as Supplementary File 1. All additional data and analysis scripts are available upon request from the corresponding author.

## Acknowledgments

The authors would like to acknowledge the members of the Pletcher laboratory for their comments on the experimental design and analysis. This research was supported by the US National Institutes of Health, National Institute on Aging (RO1AG030593 and RO1AG023166 to S.D.P); the Glenn Medical Foundation (to S.D.P.), and the NIH Cellular and Molecular Biology and Career Training in the Biology of Aging Training Grants (T32-GM007315 and T32-AG000114 to A.S.M.).

## Author Contributions

T.C., C.G., A.M., and S.P. designed the experiments. T.C., C.G., A.M., M.D., and Z.H. performed the experiments. C.G., A.M., Y.L., and S.P. analyzed the data. T.C., C.G., Y.L., A.M., and S.P. wrote the manuscript.

