## Supplemental Figures for "Perception of naturally dead conspecifics impairs health and longevity through serotonin signaling in *Drosophila*"

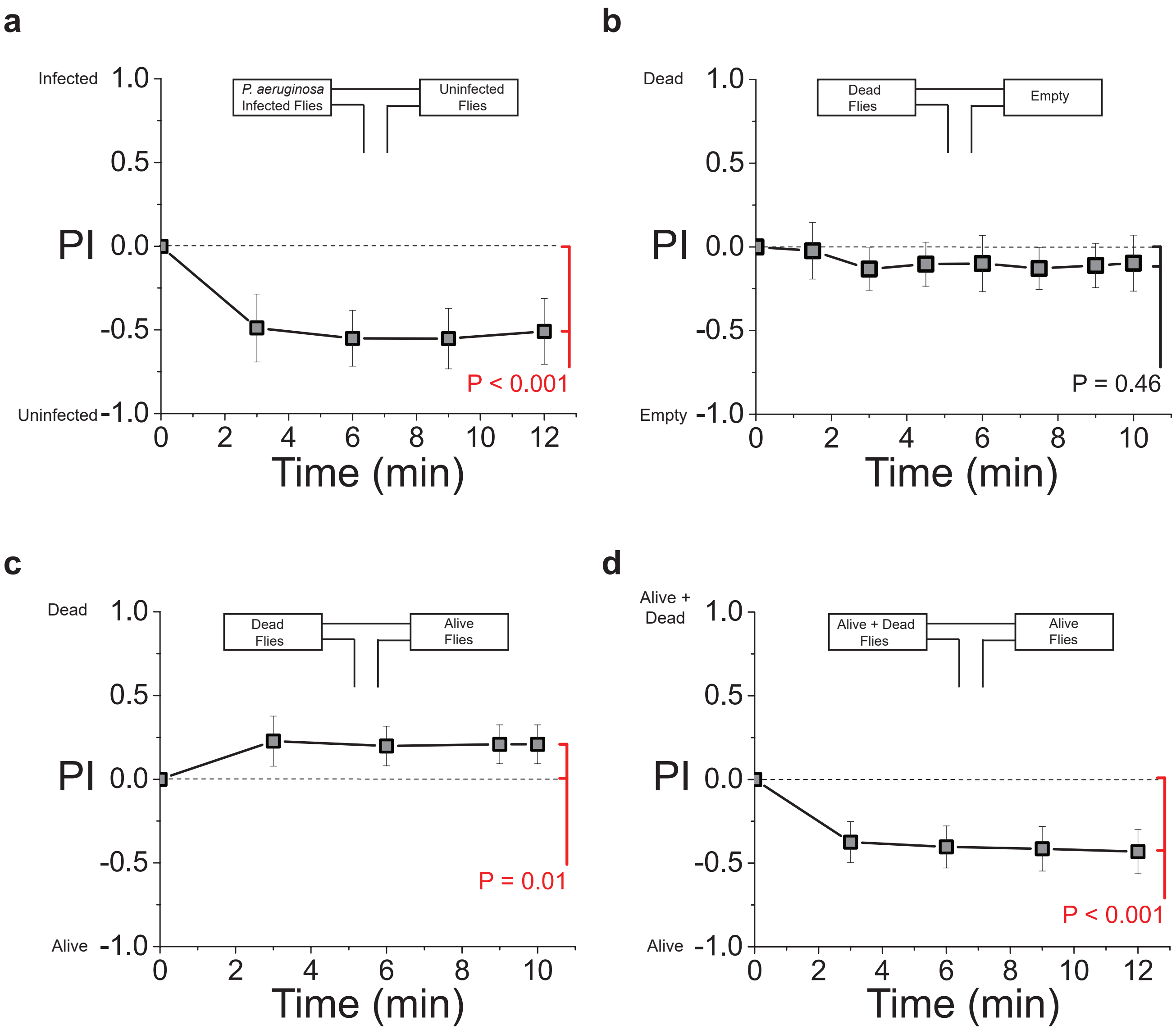


Supplemental Figure 1

**Supplement Fig. 1** Initial observations and characterization of the T-maze behavioral assay. **a** Naïve choosing flies are repelled by infected flies compared to uninfected control flies (N = 6). **b** Naïve choosing flies exhibit no preference when choosing between a chamber containing dead flies or an empty chamber (N = 20). **c** Naïve choosing flies prefer a chamber containing only dead flies over one containing only live flies (N = 4). **d** Naïve choosing flies more often choose a chamber containing only live females to a chamber that contains a mixture of live females and dead conspecifics (N = 10). All naïve choosing flies were females from the Canton-S strain. Each T-maze sample tests 20 flies.

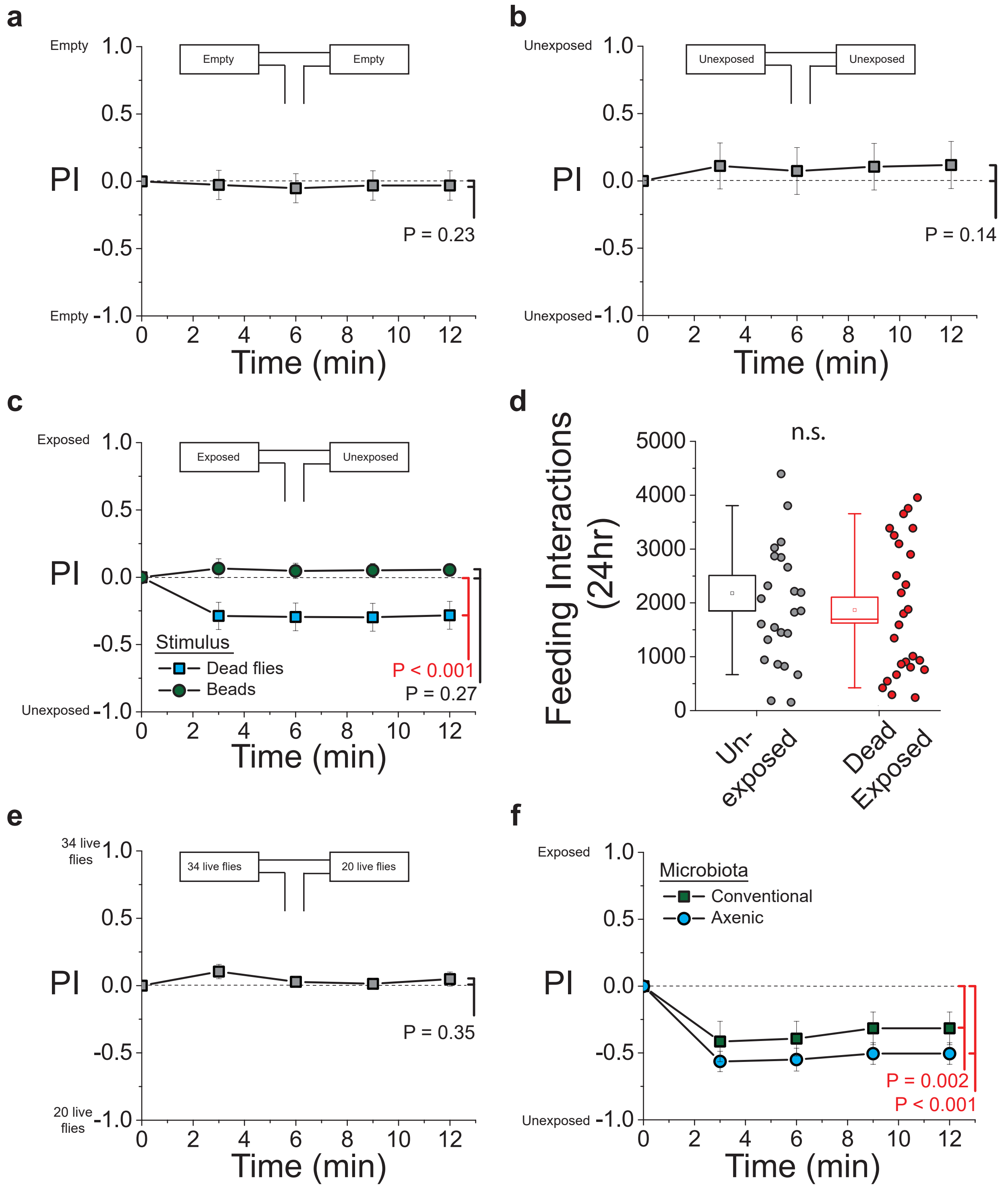


Supplemental Figure 2

**Supplement Fig. 2** Further characterization of the T-maze behavioral assay and feeding behavior. **a** Naïve choosing flies exhibit no behavioral preference when both arms of the T-maze are empty (N = 10). **b** The presence of equal numbers of live female flies in both arms of the T-maze does not result in behavioral preference (N = 8). **c** Naïve choosing flies show no preference between experimental flies that were exposed to mock dead flies (small black beads) for 48 hours relative to unexposed flies (N = 15). **d** Feeding behavior is not affected by 48 hours exposure to dead conspecifics. Each data point represents the number of feeding interactions with the food for an individual fly (N= 25 for control flies and N = 26 for exposed flies, P = 0.45). **e** Naïve choosing flies exhibit no preference between arms with different densities of live flies (34 female flies vs 20 female flies; N = 10). **f** Similar levels of aversion were induced when axenic flies were used as both exposed and dead flies as seen when using conventionally reared animals (N = 10 for both treatments). All naïve choosing flies were females from the Canton-S strain.

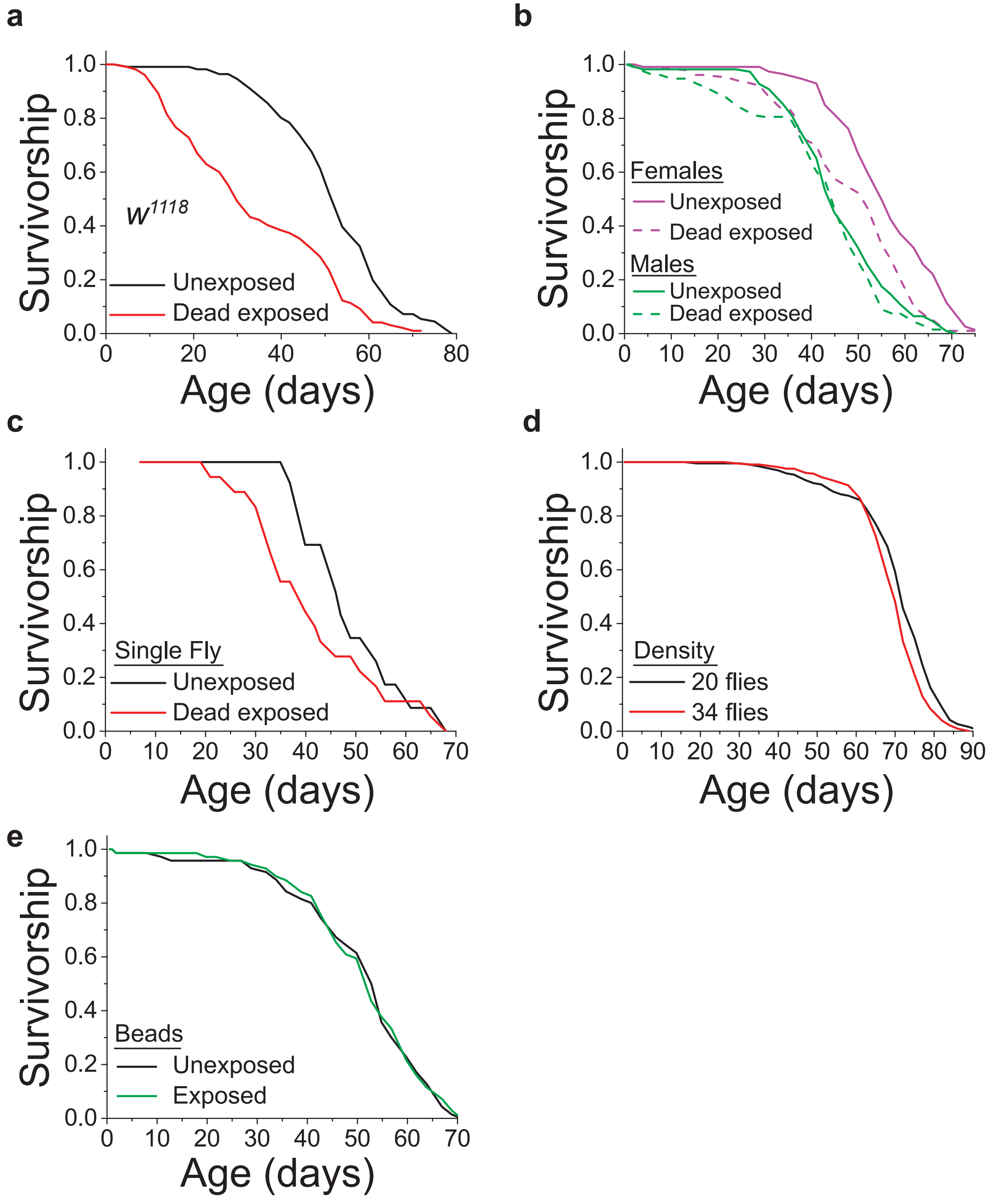


Supplemental Figure 3

**Supplement Fig. 3** The effects of death perception on lifespan with regard to genetic background, gender, and density. **a** Exposure to dead conspecifics shortens lifespan in the $w^{1118}$ genetic background, (N = 111-113 per treatment, P < 0.001). **b** Canton-S female flies exposed to dead flies live significantly shorter compared to control (unexposed) animals, whereas Canton-S male flies exposed to dead females live similarly to controls (N = 85-113 per gender/treatment, P < 0.001 for females and P = 0.33 for males). **c** Single Canton-S female flies exposed to 8 dead female flies repeatedly tend to live in shorter, although the small sample size limits statistical power (N = 16-19 per treatment, P = 0.22). **d** Increased density of live flies has a small effect on lifespan (N = 195 flies for the 20 flies/vial treatment, N = 325 flies for the 34 flies/vial treatment, P < 0.001). **e** Lifespan is unaffected when Canton-S female flies are aged in the presence of mock dead flies (14 fly-sized black beads, N = 71 for control and 70 for the beads exposed, P = 0.90).

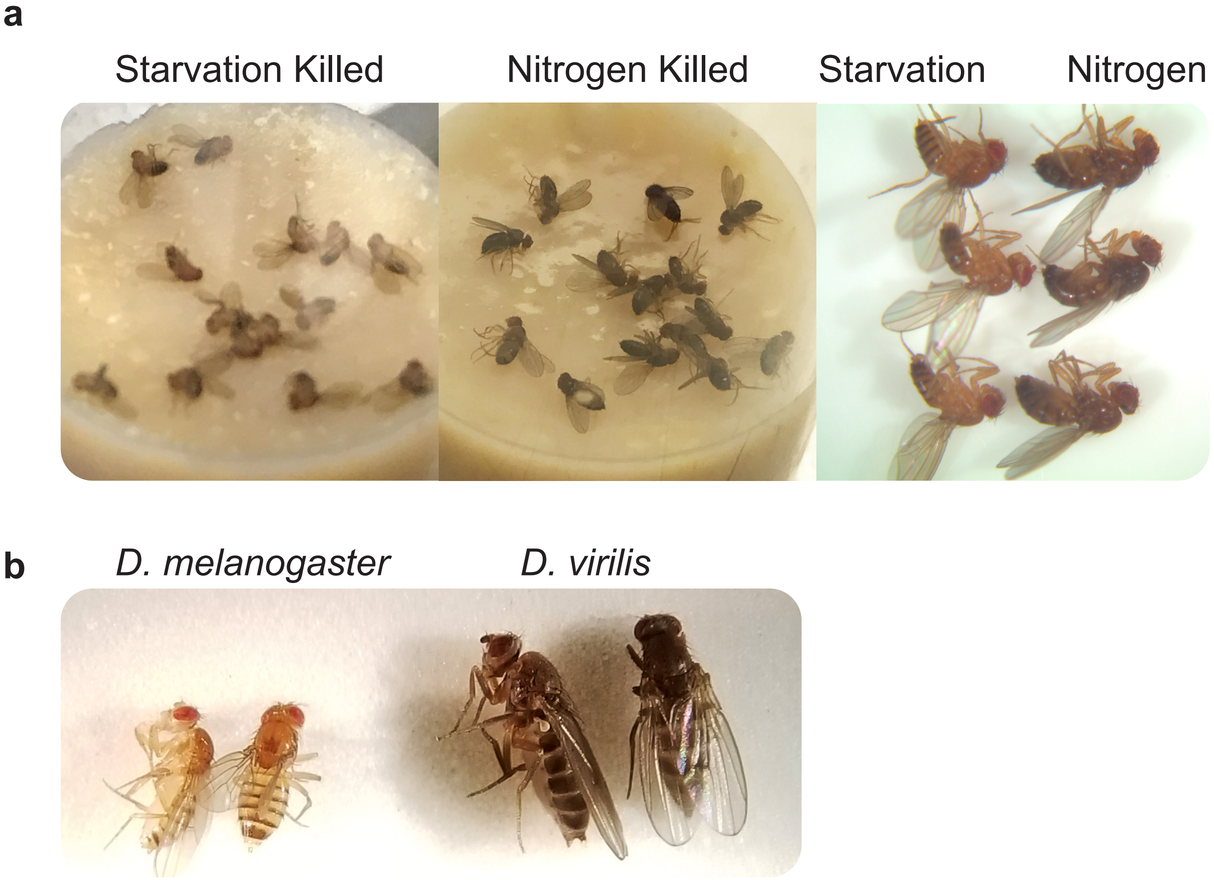


Supplemental Figure 4

**Supplemental Fig. 4** Images comparing dead flies demonstrate significant visual differences. **a** *D. melanogaster* that are killed via starvation are visually lighter in color compared to flies killed by immersion in liquid nitrogen. Both groups sat at room temperature for 2 days after dying prior to this picture being taken and were treated in exactly the same manner as those dead flies that were introduced into our experiments. **b** *D. melanogaster* are visually lighter in color and much smaller than *D. virilis*.

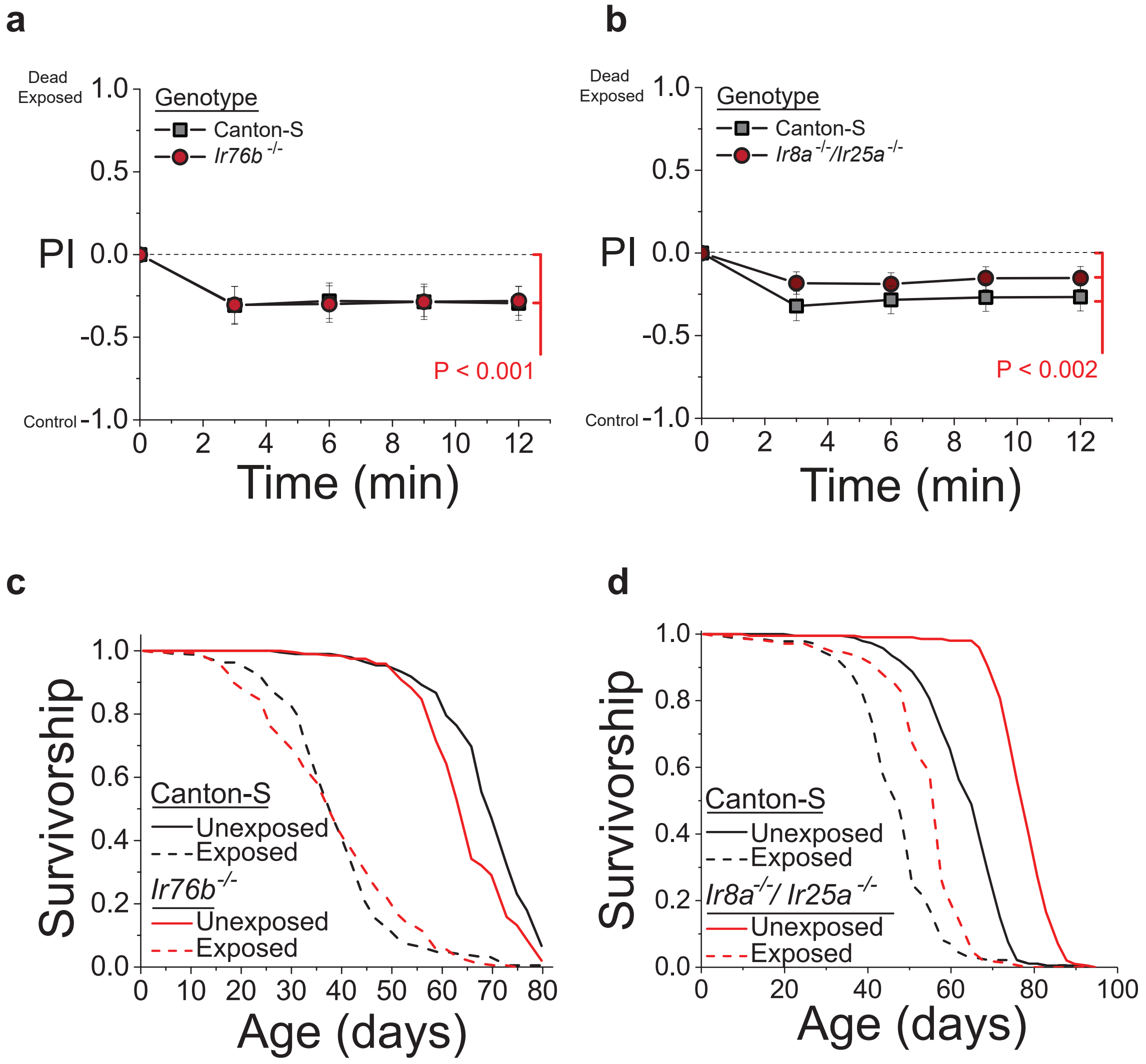


Supplemental Figure 5

**Supplemental Fig. 5** The effect of sensory manipulations on death perception part 1. **a** *Ir76b*$^{-\backslash-}$ mutant flies retained their aversive characteristics to naïve choosing flies (N = 9 for control and 10 for *Ir76b*$^{-/-}$ mutants). **b** *Ir8a*$^{-/-}$/*Ir25a* $^{-/-}$ mutant flies retained their aversive characteristics to naïve choosing flies (N = 16 for control and 17 for *Ir8a*$^{-/-}$/*Ir25a* $^{-/-}$ mutants). **c** *Ir76b*$^{-/-}$ mutant flies show a significant effect of death exposure on lifespan, comparable to controls (N = 183-197 per genotype/treatment, $P < 0.001$ for exposed vs. unexposed in both control and *Ir76b* $^{-\backslash-}$ mutants). **d** *Ir8a*$^{-/-}$/*Ir25a* $^{-/-}$ mutant flies show a significant effect of death exposure on lifespan, comparable to controls (N = 184-209 per genotype/treatment, $P < 0.001$ for exposed vs. unexposed in both control and *Ir8a*$^{-/-}$/*Ir25a* $^{-/-}$ mutants).

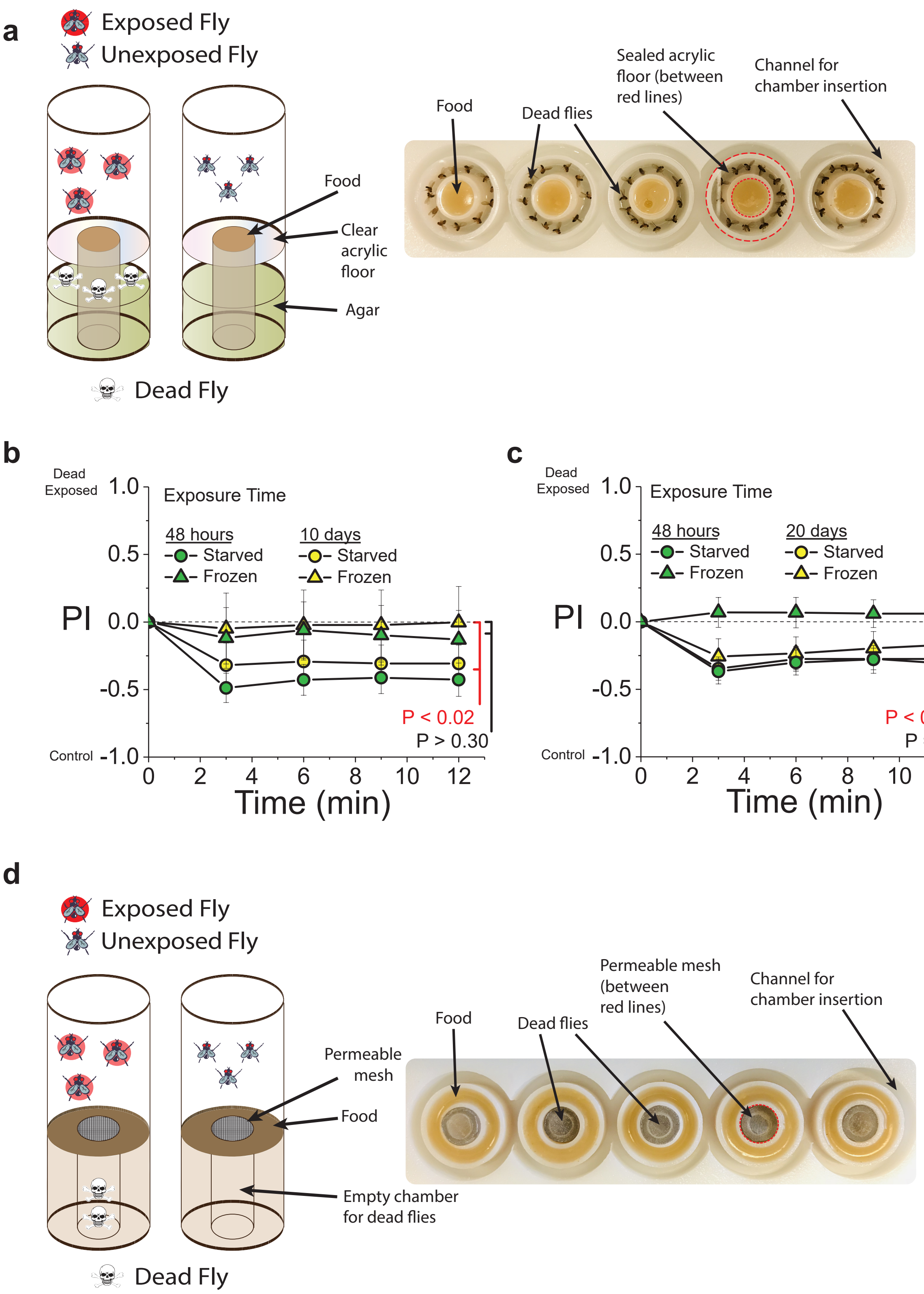


### Supplemental Figure 6

**Supplemental Fig. 6** The effect of sensory manipulations on death perception part 2. **a** Diagram of the visual exposure chamber. Flies have access to the food and can see dead flies, but cannot interact or smell the dead flies. **b** Flies that are quickly killed by immersion in liquid nitrogen do not induce avoidance behaviors when used for short term (48 hours) or medium term (10 day) exposures. Flies that are exposed to dead flies via starvation show aversion at both short and medium term exposures (N = 5 for each treatment, P = 0.30 for 48 hour dead frozen fly exposure, P = 0.0016 for 48 hour dead starved fly exposure, P = 0.42 for 10 day dead frozen fly exposure, and P = 0.016 for 10 day dead starved fly exposure). **c** Flies that are quickly killed by immersion in liquid nitrogen induced avoidance behaviors when used for long term (20 day) exposures (N = 10 for each treatment, P = 0.44 for 48 hour dead frozen fly exposure, P < 0.001 for 48 hour dead starved fly exposure, P = 0.008 for 20 day dead frozen fly exposure, and P = 0.002 for 20 day dead starved fly exposure). **d** Diagram of the odor exposure chamber. Flies have access to the food as well as odors from dead flies, but cannot interact with the dead flies. All of the flies used in these experiments were females from the Canton-S strain. Each T-maze sample tests 20 flies.

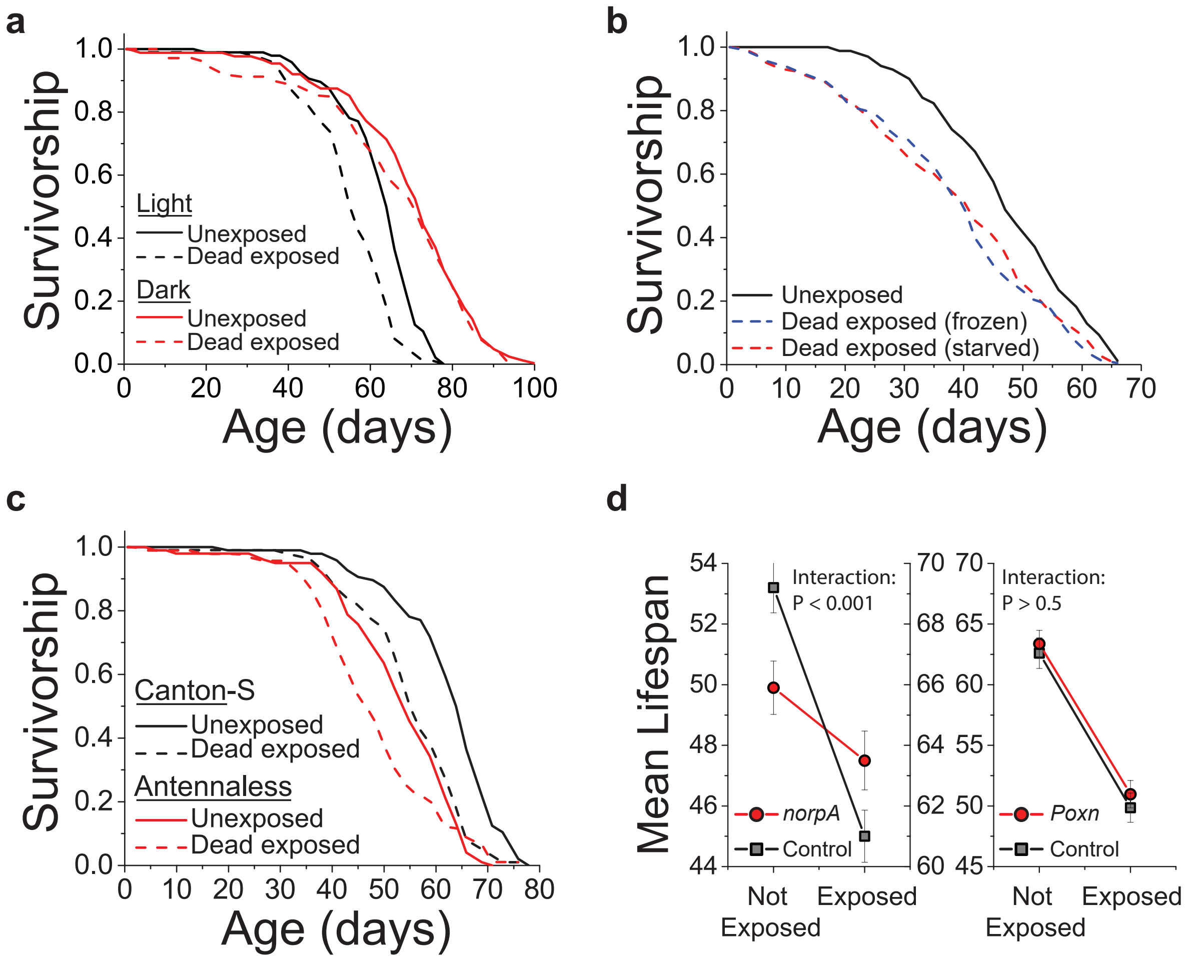


Supplemental Figure 7

**Supplemental Fig. 7** The effect of sensory manipulations on death perception part 3. **a** This experiment represents a replicate of that presented in Fig. 4a showing that when flies were exposed in the dark, dead animals failed to influence lifespan (N = 82 for dark dead exposed, 88 for dark unexposed, 101 for light dead exposed, and 96 for light unexposed, $P < 0.001$ for light and $P = 0.49$ for dark, $P < 0.001$ for the interaction between light and exposure via Cox Regression) **b** We see similar effects on lifespan when flies are exposed long-term to either dead flies that were killed via starvation or dead flies that were killed via immersion to liquid nitrogen (N = 183 for starvation-dead exposed, 201 for liquid nitrogen-dead exposed, and 200 for unexposed, $P < 0.001$ for each dead exposed treatment compared to the unexposed). **c** Canton-S flies with their antenna surgically removed maintained a reduced effect of death exposure on lifespan compared to control flies (N = 93-101 for each treatment, $P = 0.06$ for antennaless and $P < 0.001$ for control). **d** The change in mean lifespan when comparing exposed vs. unexposed cohorts of *norpA* flies is significantly decreased when compared to a similar measurement in control flies. There is no significant mean lifespan interaction when comparing exposed vs. unexposed cohorts of control or *Poxn* mutant flies.

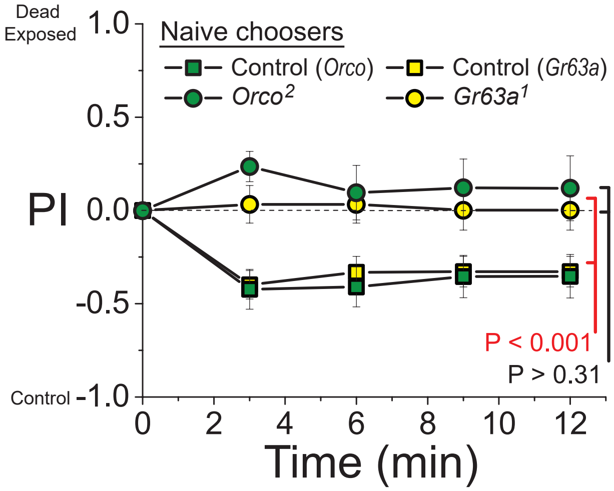


Supplemental Figure 8

**Supplemental Fig. 8** Olfaction is required in naïve choosing flies to detect the aversive cues emitted by flies exposed to dead conspecifics. In this experiment, female *Orco*$^{2}$ or *Gr63a*$^{1}$ mutants were used as the naïve choosers, while female Canton S were used as the exposed or unexposed flies in each arm of the T maze (N = 6 for *Orco* control, N = 7 for O*rco*$^{2}$, N = 10 for *Gr63a* control, and N = 10 for *Gr63a*$^{1}$). $P < 0.001$ for both controls and $P > 0.31$ for both mutants. Each T-maze sample tests 20 flies.

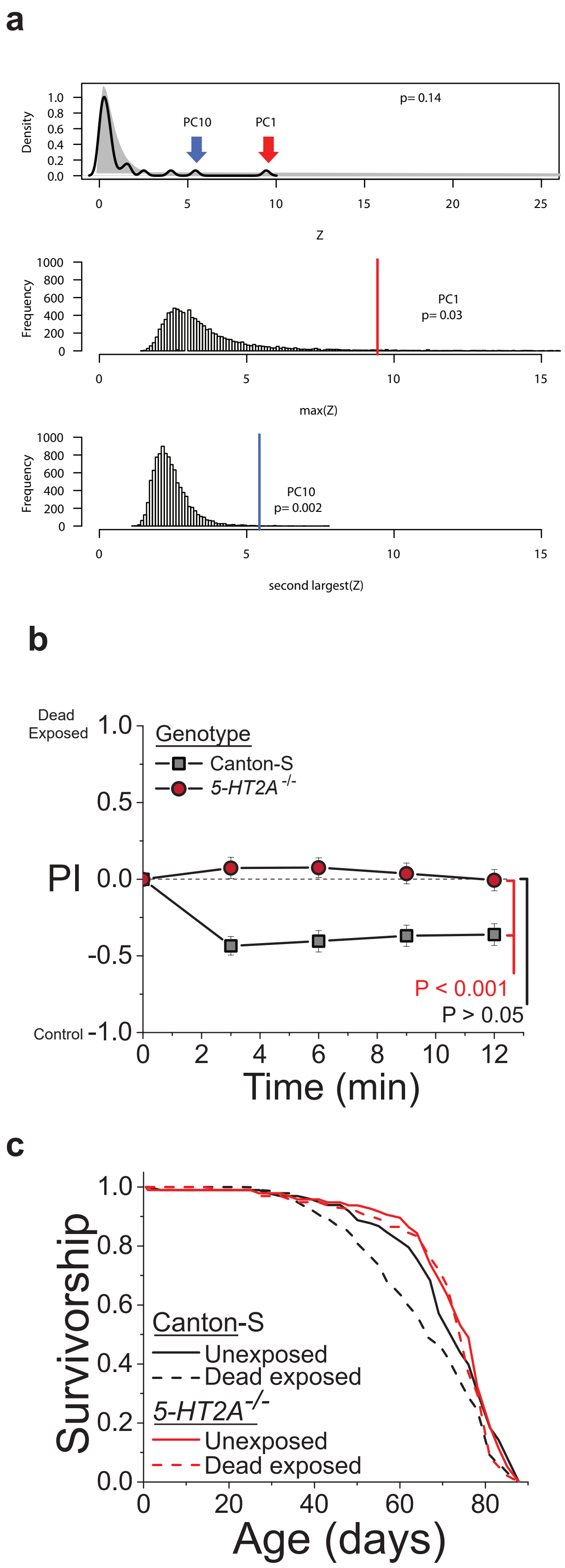


Supplemental Figure 9

**Supplementary Fig. 9** Details of the principle component analysis and effect of death perception on serotonin *5-HT2A*$^{-/-}$ mutants. **a** Principle component randomization results. Observed (black) and randomized (grey) distributions of the ability of individual principal components to effectively distinguish the neurometabolomics signature of flies exposed to conspecific dead. No statistical difference was observed between these two distributions (P = 0.14, Kolmogorov-Smirnov Test). This analysis identified two PCs, PC1 and PC10, that exhibited statistically significant ability to distinguish groups. Shown are the frequency plots and P-values for both PCs. **b** Replicate T-maze behavioral experiment showing that loss of *5-HT2A*$^{-/-}$ protected flies from the consequences of death perception on aversive cues detected by naïve choosing flies (N = 16 for each treatment). **c** Replicate survival experiment showing that *5-HT2A*$^{-/-}$ flies are protected from the consequences of death perception on lifespan (N = 94-98 per genotype/treatment, P = 0.002 for Canton-S control flies and P = 0.22 for *5-HT2A*$^{-/-}$ mutants).
