## Supplemental Table 1 for "Perception of naturally dead conspecifics impairs health and longevity through serotonin signaling in *Drosophila*"

| **Supplementary Table 1: Summary of death exposure effects on lifespan of flies fed drugs.** Values represent mean lifespan (SEM) in days for each cohort. Interaction P-values represent the results of a randomized ANOVA as described in the Methods. Note that the significant effect of ondansetron reflects an enhancement of the response, not an abrogation. | | | | | | | |
| --- | --- | --- | --- | --- | --- | --- | --- |
|  | **Vehicle (DMSO)** | | | **Drug** | | |  |
| **Drug** | **Unexposed** | **Exposed** | **% Change** | **Unexposed** | **Exposed** | **% Change** | **P** |
| Fluvoxamine | 55.69 (0.86) | 50.24 (1.13) | 10% | 56.42 (0.78) | 51.88 (0.92) | 8% | 0.35 |
| 3-Iodo-L-Tyrosine | 61.65 (1.35) | 51.93 (1.76) | 16% | 61.58 (1.36) | 49.48 (1.74) | 20% | 0.35 |
| Thiostrepton | 63.59 (1.41) | 50.77 (1.29) | 20% | 67.13 (1.38) | 53.26 (1.20) | 21% | 0.75 |
| Propantheline Bromide | 63.59 (1.41) | 50.77 (1.29) | 20% | 67.31 (1.36) | 50.47 (1.19) | 25% | 0.10 |
| Epinastine | 63.59 (1.41) | 50.77 (1.29) | 20% | 66.28 (1.24) | 50.54 (1.65) | 24% | 0.21 |
| Ondansetron | 63.59 (1.41) | 50.77 (1.29) | 20% | 70.36 (1.01) | 49.10 (1.18) | 30% | <0.001 |
| 3-Iodo-L-Tyrosine | 61.65 (1.35) | 51.93 (1.76) | 16% | 61.58 (1.36) | 49.48 (1.74) | 20% | 0.35 |
